# LIQA: Long-read Isoform Quantification and Analysis

**DOI:** 10.1101/2020.09.09.289793

**Authors:** Yu Hu, Li Fang, Xuelian Chen, Jiang F. Zhong, Mingyao Li, Kai Wang

## Abstract

Long-read RNA sequencing (RNA-seq) technologies have made it possible to sequence full-length transcripts, facilitating the exploration of isoform-specific gene expression (isoform relative abundance and isoform-level TPM) over conventional short-read RNA-seq. However, long-read RNA-seq suffers from high per-base error rate, presence of chimeric reads or alternative alignments, and other biases, which require different analysis methods than short-read RNA-seq. Here we present LIQA (<u>L</u>ong-read Isoform <u>Q</u>uantification and <u>A</u>nalysis), an Expectation-Maximization based statistical method to quantify isoform expression and detect differential alternative splicing (DAS) events using long-read RNA-seq data. Rather than summarizing isoform-specific read counts directly as done in short-read methods, LIQA incorporates base-pair quality score and isoform-specific read length information to assign different weights across reads, which reflects alignment confidence. Moreover, LIQA can detect DAS events between conditions using isoform usage estimates. We evaluated LIQA’s performance on simulated data and demonstrated that it outperforms other approaches in characterizing isoforms with low read coverage and in detecting DAS events between two groups. We also generated one direct mRNA sequencing dataset and one cDNA sequencing dataset using the Oxford Nanopore long-read platform, both with paired short-read RNA-seq data and qPCR data on selected genes, and we demonstrated that LIQA performs well in isoform discovery and quantification. Finally, we evaluated LIQA on a PacBio dataset on esophageal squamous epithelial cells, and demonstrated that LIQA recovered DAS events that failed to be detected in short-read data. In summary, LIQA leverages the power of long-read RNA-seq and achieves higher accuracy in estimating isoform abundance than existing approaches, especially for isoforms with low coverage and biased read distribution. LIQA is freely available at https://github.com/WGLab/LIQA.

## Introduction

RNA splicing is a major mechanism for generating transcriptomic variations, and mis-regulation of splicing is associated with a large array of human diseases caused by hereditary and somatic mutations[1-5]. Over the past decade, RNA sequencing (RNA-seq) has revolutionized transcriptomics studies and facilitated the characterization and understanding of transcriptomic variations in an unbiased fashion. With RNA-seq, we can quantitatively measure isoform-specific gene expression, discover novel and unique transcript isoform signature and detect differential alternative splicing (DAS) events[6-8]. Until now, short-read RNA-seq has been the method of choice for transcriptomics studies due to its high coverage and single nucleotide resolution[8]. However, due to limited read length, it is difficult to accurately characterize transcripts using short reads, as 81% of isoforms have length greater than 500bp in the GENCODE annotation (median = 1,543 bp and mean = 2,121 bp). This fragmented sequencing of the RNA/cDNA molecules results in biases and has become a barrier for short reads to be correctly mapped to the reference genome, which is crucial for gene or isoform expression estimation and novel or unique isoform detection. As a consequence, transcriptome profiling using short-read RNA-seq is highly biased by read coverage heterogeneity across isoforms. To tackle these challenges, a number of computational tools, including RSEM[9], eXpress[10], TIGAR2[11], Salmon[12], Sailfish[13], Kallisto[14], Cufflinks[15], CEM[16], PennSeq[17], IsoEM[18], and RD[19], have been developed to quantify isoform expression from short-read RNA-seq data, but different bias correction algorithms can result in conflicting estimates[17]. Overall, quantifying isoform expression using fragmented short reads remains challenging, especially at complex gene loci[20, 21].

In recent years, long-read RNA sequencing has gained popularity due to its potential to overcome the above-mentioned issues when compared to short-read RNA-seq[22, 23]. Previous studies have utilized both single-molecule long-read PacBio Iso-Seq and synthetic long-read MOLECULO methods[24-27]. For Oxford Nanopore sequencing, there are two types of RNA-seq technologies: direct mRNA sequencing and cDNA sequencing. Recently, the Oxford Nanopore Technologies (ONT) MinION has been used to analyze both full-length cDNA samples and mRNA samples derived from tissue cells[28]. Nanopore sequencing is able to generate reads as long as 2Mbp, which allows a large portion or the entire mRNA or cDNA molecule to be sequenced. Compared to short-reads, this advantage of long-reads greatly facilitates rare isoform discovery, isoform expression quantification and DAS event detection.

However, there are still a few unique challenges to analyze long-read RNA-seq data because existing methods developed for Illumina short-read RNA-seq do not have optimal performance when directly used on long-read RNA-seq. This is because that parametric bias correction of short-read approaches require high read coverage and isoform-read assignment is not robust to small range misalignment from long-read data[16, 18, 19, 29]. Methods designed specifically for isoform expression estimation in long-read RNA-seq have only emerged recently. For example, Byrne *et al*[30] demonstrated the feasibility of quantifying complex isoform expression using Nanopore RNA-seq data. Tang *et al*[31] characterized mutated gene *SF3B1* at isoform level in chronic lymphocytic leukemia cells by leveraging full-length transcript sequencing data generated by Nanopore. While long-read RNA-seq has great potential, the isoform quantification accuracy is still constrained by high error rates and sequencing biases[32], which has yet to be thoroughly accounted for. Specifically, high sequencing error rates (∼15%) of Nanopore data can result in misalignment of sequencing reads, but current methods assume equal weight for each single molecule read without accounting for error rate differences when estimating isoform expression. This may complicate isoform usage quantification. In addition, potential read coverage biases are not explicitly taken into account by existing long-read transcriptomic tools[32]. In Nanopore direct RNA sequencing protocol, pore block and fragmentation can result in truncated reads, leading to biased coverage towards the 3’ end of a transcript[32]. These biases are also shown in data sequenced from cDNA. In the presence of such biases, the accuracy of isoform expression quantification inference can be severely affected, leading to over estimation of expression for isoforms with short length.

In this article, we present LIQA, a statistical method that allows each read to have its own weight when quantifying isoform expression. Rather than counting single molecule reads equally, we give a different weight to each read to account for read-specific error rate and alignment bias at the gene (Figure 1). To evaluate the performance of LIQA, we simulated long data with known ground truth and also sequenced two real samples using Oxford Nanopore sequencing. Our results demonstrate that LIQA is an accurate approach for isoform expression quantification accounting for read coverage bias and high error rate of long-read data.

**Figure 1.**
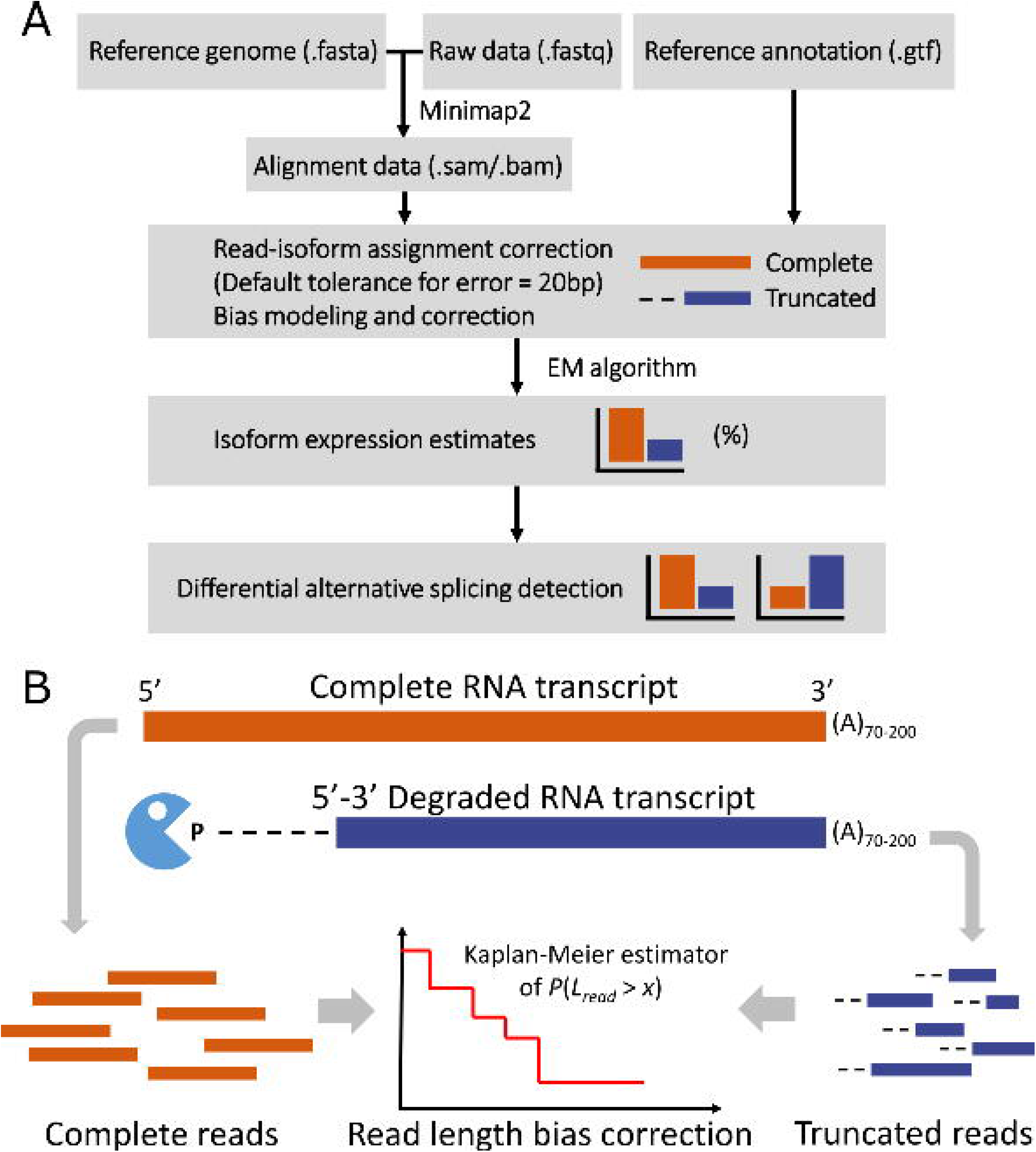
Framework of LIQA. (A) The flowchart to illustrate how LIQA works. The inputs to LIQA are long-read RNA-seq data and isoform annotation file. LIQA models observed splicing information, high error rate of data and read length bias. The output of LIQA are isoform expression estimates and detected DAS events. (B) Quantification of potential 3’ bias of long-read RNA-seq data. Complete and degraded RNA transcript are indicated by orange and blue. Complete (orange) and truncated (blue) long reads are jointly modeled to correct read length bias by estimating read length distribution.

## Results

### Overview of LIQA

Figure 1 shows the workflow of LIQA and highlights the read length bias correction step. LIQA requires aligned long-read RNA-seq files in BAM or SAM format and isoform annotation file as input. For estimation steps, LIQA first feeds read alignment information to a complete likelihood function and corrects biases for each long read by accounting for quality score and read coverage bias. Second, given that isoform origins are unobserved for some reads, an Expectation Maximization (EM)-algorithm is utilized to achieve the optimal solution of isoform relative abundance estimation. The output values of LIQA are isoform expression estimates. Moreover, LIQA can further detect DAS events given estimated isoform expression values.

To evaluate the performance of LIQA, we compared it with existing long-read based quantification algorithms, including FLAIR [31], Mandalorion [30], TALON [33] and the Oxford Nanopore Pipeline (ONP, https://github.com/nanoporetech/pipeline-transcriptome-de). These methods use long-read RNA-seq data to detect novel isoforms and quantify transcript expression by counting the number of reads, which give equal weight for each read. To make the comparisons fair, we ran LIQA, FLAIR. TALON, Mandalorion and ONP in quantification mode only with isoform annotation information provided by GENCODE. We benchmarked the performance of each method on both simulated and real data. In addition, we simulated more data to evaluate the performance of LIQA in detecting DAS events between conditions.

### Nanopore RNA-seq data simulation

We conducted a simulation study to evaluate the performance of LIQA and compared it with other state-of-the-art algorithms for isoform expression estimation and DAS detection based on GENCODE v24 annotation. To simulate a realistic dataset with known ground truth, we used NanoSim[34] to generate the ONT RNA-seq data. NanoSim is a fast and scalable read simulator that captures the technology-specific features of ONT data, and allows for adjustment upon improvement of Nanopore sequencing technology. This simulator first characterizes Nanopore reads and models features of the library preparation protocols *in silico* for read simulation. The human genome sequence (GRCh38), transcriptome sequence and GTF annotation file were downloaded from GENCODE. To make the simulated data close to real studies, we assigned abundance values for each isoform obtained from a real human eye RNA-seq dataset. Using NanoSim, we generated 5 million (5M) Nanopore reads. To evaluate the impact of sequencing depth on isoform expression quantification, we down-sampled 3 million (3M), 1 million (1M) and 0.5 million (0.5M) reads for the simulated data, respectively. These reads were aligned to the reference genome using minimap2 [35]. Then, we selected genes with 2 or more isoforms (67.2%) to evaluate the performance of LIQA in isoform expression quantification. For each isoform, we compared it with Mandalorion, FLAIR, TALON and ONP. All methods were run with the same set of simulated aligned data in BAM format as input.

The characteristics of the simulated data are shown in Figure 2(A) and Supplementary Figure 1(A). The median lengths of ONT reads in the 0.5M, 1M, 3M and 5M datasets are 896, 922, 1,010 and 993 base pairs, respectively. Among the evaluated genes with multiple isoforms (67.2%) based on GENCODE annotation, 13% have two isoforms, 14% have three isoforms and 73% have four or more isoforms. The simulated isoforms have a wide range of relative abundance (interquartile range = (0.002, 0.75), median = 0.041). In addition, by training the statistical model of NanoSim with a real long-read RNA-seq dataset, the coverage plots of the simulated data capture the features of real ONT RNA-seq data, demonstrating 3’ bias (Supplementary Figure 1(B)). These simulated data thus provide an ideal basis to evaluate the performance of LIQA as the ground truth is known.

**Figure 2.**
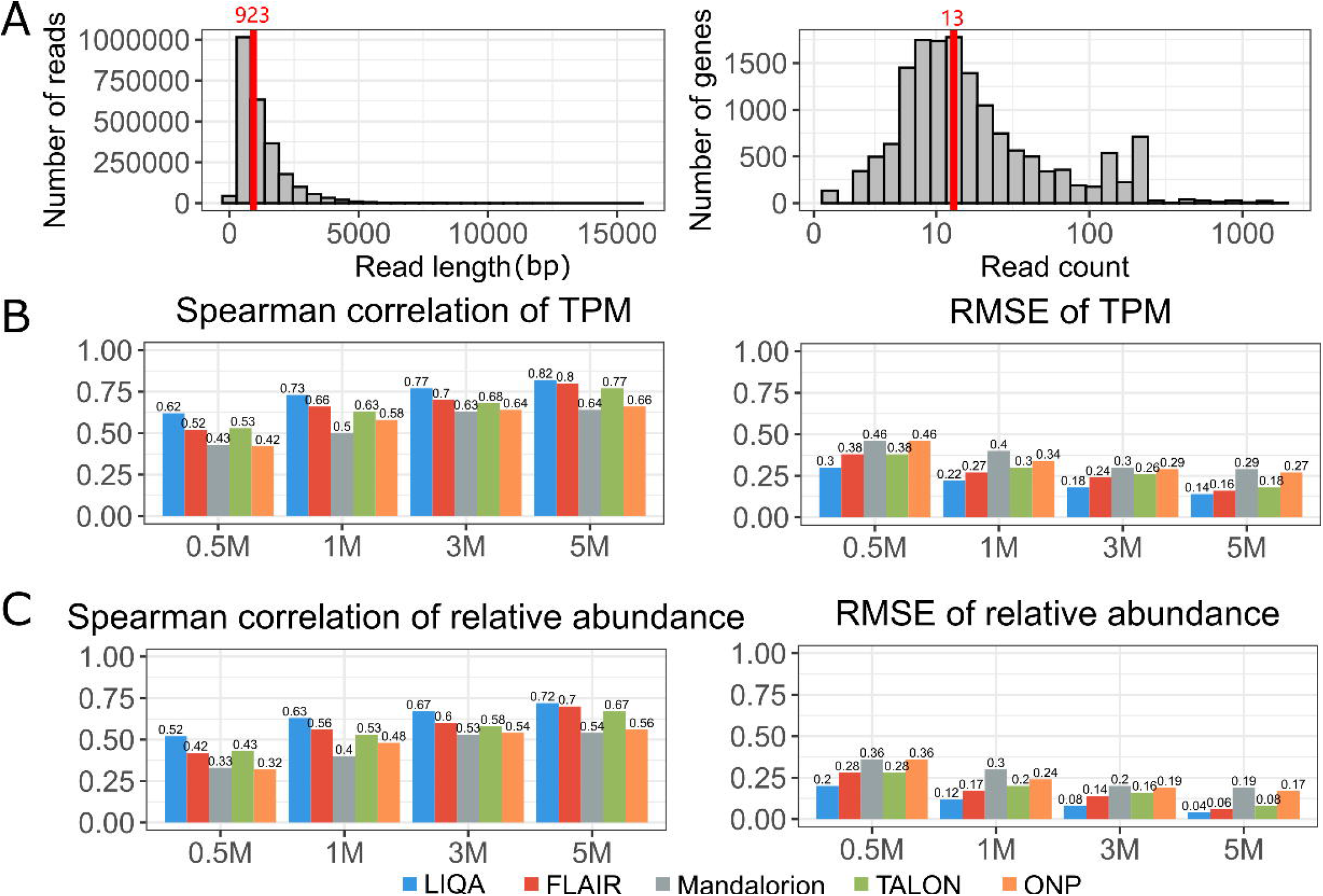
Simulation study results. (A) Characteristics of simulated data with 5M. Read length distribution (left) and read count distribution by genes in log-scale (right). (B-C) Summary statistics between true and estimated isoform expressions using LIQA (blue), FLAIR (red), Mandalorion (gray), TALON (green) and Oxford Nanopore Pipeline (ONP) (yellow) at different read coverages. (B) Spearman’s correlations (left) and RMSE (right) between estimated and true TPM. (C) Spearman’s correlations (left) and RMSE (right) between estimated and true relative abundance.

### Gene or Isoform expression quantification accuracy

For each simulated dataset, we computed a set of measurements to evaluate the estimation accuracy of each method. First, we measured the similarity between the estimated isoform relative abundance and the ground truth by calculating the Spearman’s correlation. Second, we measured the estimation accuracy by calculating the root mean squared error (RMSE), defined as 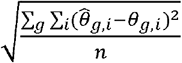, where the summation is taken over all genes and all isoforms within each gene and *n* is the total number of isoforms across all genes. Both statistics were computed at three levels: global gene expression, global isoform expression, and within-gene isoform relative abundances.

Figure 2(B)(C) (Supplementary Figure 2) show the summary statistics between estimated and true values of global isoform expression (global isoform expression and isoform relative abundances) at different read coverages. Spearman’s correlation and RMSE were calculated for all three methods. LIQA, FLAIR and TALON have higher Spearman’s correlation than Mandalorion and ONP for simulated datasets with low sequencing depth (1M, 0.5M) (Supplementary Table 2). For simulated data with high sequencing depth (3M, 5M), LIQA has similar estimation accuracy with FLAIR and TALON based on Spearman’s correlation (Supplementary Table 2). Figure 2(C) gives summary statistics of relative abundance estimates for the five methods. For relative abundance estimation, LIQA outperforms FLAIR and TALON with 6.6% and 7% lower RMSE on average, respectively. Comparison results at gene level reveal similar pattern (Supplementary Figure 4, Supplementary Table 5). The improved performance of LIQA is likely due to its use of the EM-algorithm, which assigns unequal weight to each read to better account for mapping uncertainty and read mapping bias (Figure 3(C)(D) and Supplementary Table 6, Supplementary Table 7). In contrast, FLAIR, TALON and Mandalorion provide discrete estimations by directly counting the number of reads aligned to each corresponding gene or isoform. Due to the limited read coverage of ONT RNA-seq data, it is not surprising that they yield lower estimation accuracy.

**Figure 3.**
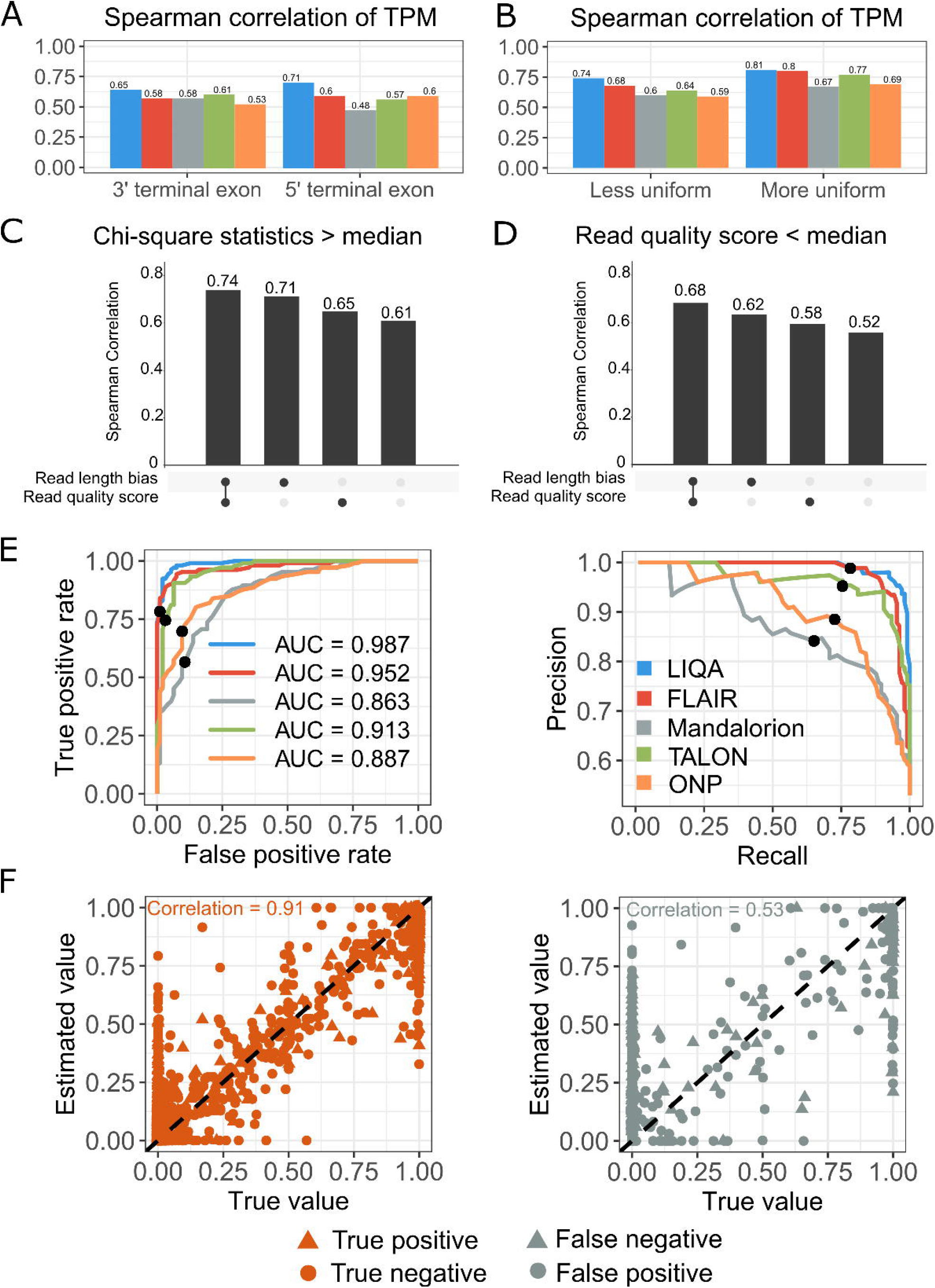
Evaluation of robustness to read length bias and low read quality (A-D) and comparison of DAS events detection (E-F) using LIQA (blue), FLAIR (red), Mandalorion (gray), TALON (green) and ONP (yellow). (A) Spearman’s correlations between estimated and true TPM for 3’ and 5’ terminal exons. (B) Spearman’s correlations between estimated and true TPM. Isoforms are stratified by the chi-squared goodness of fit statistic for uniformity. The left panel is for those isoforms in which the read coverage distribution is less uniform, and the right panel is for the remaining isoforms. (C-D) Spearman’s correlation comparison between different models (full model, read length model, read quality model and baseline model) based on isoforms with chi-squared statistic more than the median (C) and isoforms with average read quality score less the median (D). (E) ROC curve and precision recall curve of different methods for DAS gene detection. Threshold FDR<0.05 is highlighted using black dot. (F) Scatter plot of true and estimated isoform relative abundance from the first DAS simulation dataset. LIQA’s prediction and ground truth are marked for each isoform.

To evaluate the robustness of LIQA to a more complex isoform annotation, we analyzed 5M simulation dataset based on GENCODE v37 annotation. For major use isoforms, as shown in Supplementary Figure 4(D), LIQA still yields 7% higher spearman correlation of TPM and relative abundance estimates than second best approach FLAIR. For over-annotation, we simulated RNA-seq reads based on 66% of the GENCODE v37 annotation. We then analyzed the simulated data with various methods using 100% of the GENCODE annotation, corresponding to 50% more of the true annotation. Supplementary Figure 4(E) shows the Spearman correlation results under different degrees of over-annotation. We find that the quantification accuracy of LIQA remained nearly unchanged (1% less) as the degree of over-annotation increased. For under-annotation, we simulated RNA-seq reads based on 100% of the GENCODE v37 annotation. We then analyzed the simulated data using 50% of the GENCODE annotation, corresponding to 50% less of the true annotation. Supplementary Figure 4(E) shows the Spearman’s correlation results under different degrees of under-annotation. The estimation accuracy is 10% lower when 50% less of the true annotation was used in the analysis.

Next, we evaluated the robustness of LIQA to 3’ read coverage bias (Figure 3 and Supplementary Figure 3). First, we compared the accuracy statistics for 5’ terminal exon and 3’ terminal exon of each isoform. Isoform expression with non-uniform read coverage is more challenging to estimate because the 5’ end is less likely to be covered by sequencing reads compared to 3’ end. Figure 3(A) shows the comparison of Spearman’s correlation for five methods with 0.5M read coverage. LIQA is more accurate than the other four methods at 5’ terminal exon, especially when sequencing depth is low (Supplementary Table 4). The Spearman’s correlation coefficient of LIQA is 11% higher than the second best performing method FLAIR for 5’ terminal exons, while only 6% higher for 3’ exons. This improved performance of LIQA in terminal exons quantification is also demonstrated by RMSE values. LIQA has 8%-15% improvement of RMSE values compared to other methods. Second, we considered the chi-square statistics that measures the goodness-of-fit of coverage uniformity. Then, we divided the isoforms into two categories based on median of the corresponding measure (Chi-square statistic > median, Chi-square statistic < median) and summarized Spearman’s correlation coefficient and RMSE for each group of isoforms. For isoforms with more uniform read coverage, Spearman’s correlations of LIQA and FLAIR are close. However, despite reduced Spearman’s correlation value, LIQA is more accurate than the other four methods for isoforms with less uniform read coverage (Chi-square statistic > median) (Supplementary Table 4). The improvement of LIQA compared to FLAIR is 5% higher for these isoforms. This is likely because LIQA models potential truncated reads which result in 3’ coverage bias when quantifying isoform expression.

Moreover, we assessed the impact of modeling read length bias and read quality score on the accuracy of isoform expression estimation. Figure 3(C)(D) show the comparison of isoform estimation accuracy using different models. For isoforms with less uniform read coverage (Chi-square statistics > median), model with read length bias correction has 9% (Full model vs read quality model) and 10% (Read length model vs baseline model) higher Spearman’s correlation. For genes with less average read quality, model with read quality score has 6% higher Spearman’s correlation. Overall, isoform estimation accuracy drops noticeably when using baseline model (Supplementary Table 7). This comparison demonstrates the advantage of LIQA in handling read length bias and 3’ bias correction over other approaches.

### Differential alternative splicing (DAS) detection

Next, we evaluated the performance of LIQA in DAS detection. More ONT RNA-seq data across multiple samples (*10* cases and *10* controls) were simulated for 10 times. NanoSim generated 3 million reads based on the GENCODE annotation per sample. To make true DAS events more realistic, we sampled relative abundances of isoforms from a Dirichlet distribution with mean and variance parameters estimated from a human eye RNA-seq dataset. Similarly, these simulated data were mapped to the hg38 human reference genome using minimap2. Isoform expression and usage difference between conditions were quantified using LIQA, FLAIR, TALON, Mandalorion and ONP, respectively. We first compared the performance of DAS detection between these methods using three summary statistics. After FDR control, we measured the recalls (power) of our method by calculating the proportion of correctly predicted DAS events among true DAS events. Second, we obtained precisions by calculating the proportion of correctly predicted DAS events among predicted DAS events. Additionally, F1 score 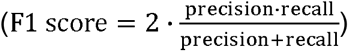 was summarized to average the precision and recall values.

As shown in Supplementary Figure 5(B), LIQA, FLAIR and TALON are more powerful than others for all three evaluation metrics. This is not surprising because Madalorion and ONP have lower accuracy in isoform expression estimation. For recall value, FLAIR (mean = 0.809, SD = 0.041) gives better and more consistent performance across 10 simulations than LIQA (mean = 0.776, SD = 0.058). However, in terms of precision value, LIQA (mean = 0.915, SD = 0.043) yields less false positives than FLAIR (mean = 0.884, SD = 0.051). LIQA, FLAIR and TALON had similar performance in detecting DAS events based on F1 score. Furthermore, we generated ROC curve and precision-recall curve to compare the performance between methods at different FDR thresholds (Figure 3(E) and Supplementary Figure 6, 7). As shown in Figure 3(E), LIQA achieved AUC = 0.94 after FDR control (FDR < 0.05). Given FDR threshold equals to 0.05, LIQA gave the best performance with precision = 0.98 and recall = 0.78. Compared to LIQA, the second-best performing method FLAIR yields 0.1%, 0.3%, 3.5% less in precision, recall and AUC respectively. In addition, we examined isoform relative abundance estimation accuracy from correct and incorrect detected DAS genes by LIQA. After FDR control, we identified that 537 out of 2,465 genes are significantly differential spliced, which 431 are true positives and 1,836 are true negatives. For these correctly predicted genes, true isoform relative abundance is highly correlated with LIQA’s estimates (Spearman’s correlation= 0.91). For false positive and negative genes, the Spearman’s correlation is 38% lower compared to true positive and negative. This is not surprising because accurate estimation of isoform expression level leverages the power of regression model in detecting DAS events.

### Application to the Universal Human Reference (UHR) RNA-seq data

As NanoSim generates ONT RNA-seq data based on trained parametric statistical model, we recognized that simulated data is hardly a full representation of reality. To evaluate the performance of LIQA in a real setting, we sequenced the Universal Human Reference sample with Nanopore Direct mRNA sequencing (Supplementary Figure 12). Then, the resulting ONT-RNA-seq data were analyzed using all five long-read based methods (LIQA, FLAIR, Mandalorion, TALON, ONP). As quantitative real time polymerase chain reaction (qRT-PCR) is considered as the most reliable technology for measuring true isoform abundance, we downloaded the qRT-PCR measurements from the MAQC project under Gene Expression Omnibus with accession number GSE5350. As part of the MAQC project, the expression levels of 894 isoforms were measured by TaqMan Gene Expression Assay based qRT-PCR. Additionally, we downloaded the UHR short-read RNA-seq data generated using the Illumina platform. This dataset was analyzed using Cufflinks[15], CEM[16], Salmon[36] and Kallisto[14] to compare the performance in isoform quantification between long reads and short reads. Specifically, we mapped ONT and Illumina sequenced reads to the reference genome using Minimap2[35] and STAR[37], respectively, and applied each quantification method to the RNA-seq data. qRT-PCR measurements were treated as gold standard to compare the performance across methods. We note that 563 of the 894 transcripts with qRT-PCR measurements are from genes with a single isoform. Estimation results from these genes were served as positive controls (Supplementary Figure 8(A)) because estimating isoform-specific expression for these single-transcript genes is trivial. To compare the performance across different methods, we considered those transcripts that are derived from genes with two or more isoforms.

To assess the accuracy between estimates and qRT-PCR measurements, we summarized similarity metrics (Spearman’s correlation and Pearson’s correlation) of the isoform abundance values in log-scale. As shown in Figure 4(A)(B), the estimation accuracy of all methods is lower than simulated data, especially for those transcripts with qRT-PCR measures close to 0. Nevertheless, we observed consistent results in terms of relative performance of different methods with simulation data. LIQA is more accurate other methods with stronger linear relationship between logarithm estimates and qRT-PCR measurements. However, many of the lowly to moderately expressed isoforms were underestimated using the other methods with their TPM values being compacted toward 0. For ONT data, the spearman’s correlation of LIQA is 0.68, whereas the corresponding values from second best method TALON is 0.57. For Illumina data, Cufflinks seems to correlate with the qRT-PCR measurements better than others (Supplementary Figure 8(B)). The main reason for the better performance of LIQA is likely due to quantifying isoform expression by accounting for isoform length bias and base quality scores. Moreover, we randomly selected 3 genes and generated sashimi plots in Figure 4(C) to show the read coverage difference between direct mRNA sequencing and Illumina data. Overall, read distribution of long-read data is less heterogeneous than short-read. Specifically, for gene *CAPNS1*, there is clearly severe 5’ degradation in Illumina data, but with full length and even coverage across the transcripts for long-read data. Terminal exons at 5’ end in red square are crucial informative regions for splicing analysis, which enable us to differentiate read origin from different isoforms. As shown in Figure 4(C), these exonic regions were captured by Nanopore reads but missed by Illumina reads, which significantly facilitates isoform expression quantification using long-read RNA-seq data. Similarly, sashimi coverage plots of other two genes showed the same pattern, which demonstrates the advantage of long-read data over short-read in alternative splicing study.

**Figure 4.**
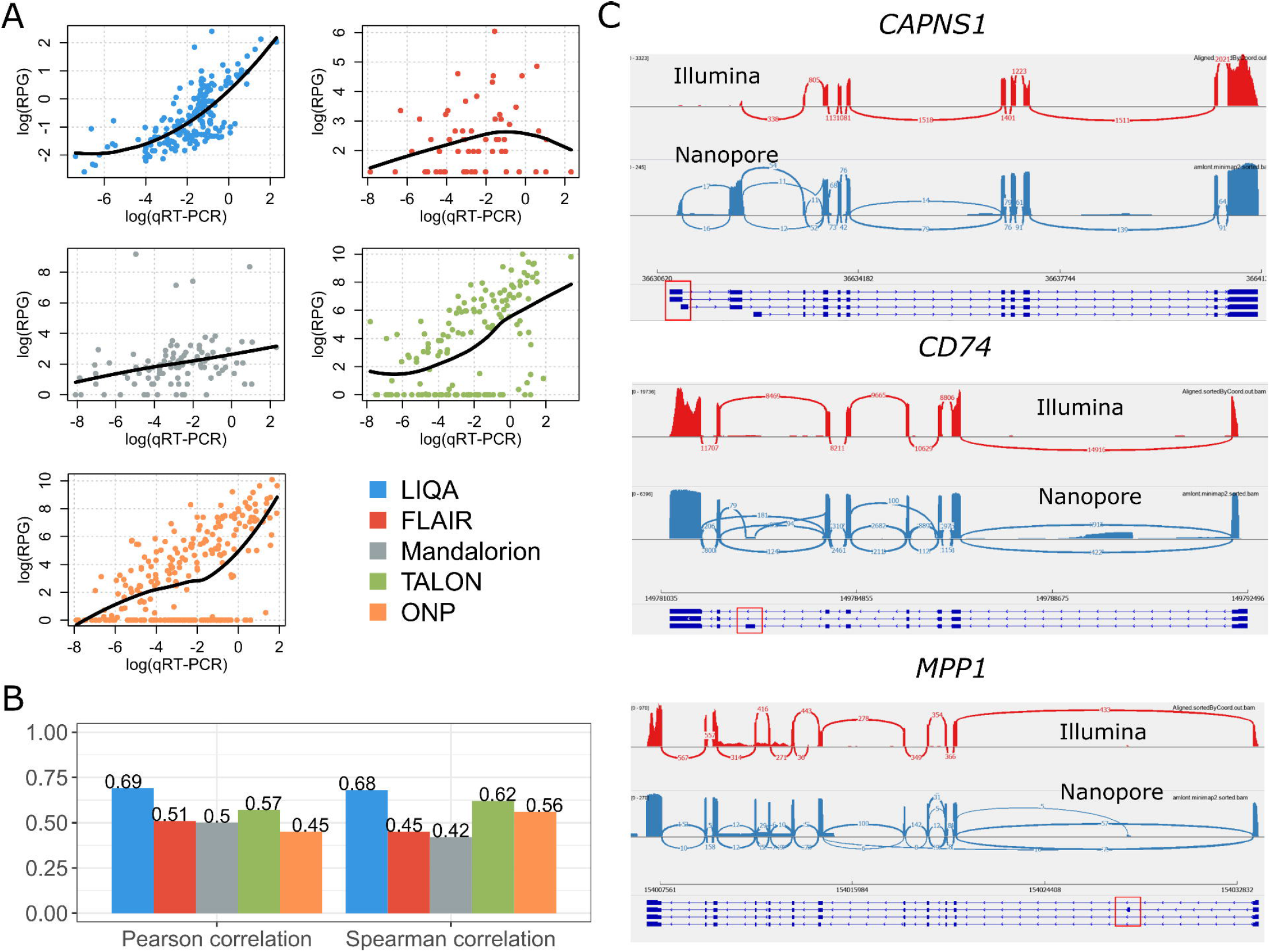
Results on UHR data generated by direct mRNA sequencing on the Nanopore platform. (A) Scatter plots of estimated isoform-specific expression versus qRT-PCR measurements in log scale. Black line represents the local polynomial regression line fitted with estimates and qRT-PCR in log scale (B) Pearson’s correlation coefficients (left) and spearman’s correlation coefficient (right) between estimated isoform-specific expression versus qRT-PCR measurements in log scale. (C) Examination of read coverage difference between Illumina and Nanopore data at 3 genes. Informative exonic regions were in red square.

Moreover, we conducted additional analysis of another long-read data on UHR with much higher coverage (5.6 million reads), generated on the PacBio sequencing platform [38]. As shown in Supplementary Figure 8(E), Pearson and Spearman’s correlations for each method are generally improved with increasing sequencing depth compared to our Nanopore-based UHR dataset (Figure 4(B)). For example, Spearman’s correlation of FLAIR is increased by 0.28 (from 0.45 to 0.73), whereas the corresponding values of LIQA is increased by 0.11. Nevertheless, LIQA still has the best performance among all methods. Based on this real dataset with increasing sequencing depth, we found that LIQA is more robust to low read coverage compared to FLAIR, which performs well when sequencing depth is high. These observations from these two real UHR datasets are consistent with the simulation based datasets with different sequencing depths (0.5M, 1M, 3M, 5M).

### Application to Nanopore cDNA sequencing data on a patient with acute myeloid leukemia

AML is a type of blood cancer where abnormal myeloblasts are made by bone marrow[39]. In this study, we sequenced peripheral blood from an acute myeloid leukemia (AML) patient using GridION Nanopore sequencer with Guppy basecalling (https://denbi-nanopore-training-course.readthedocs.io/en/latest/basecalling/basecalling.html). In total, we generated 8,061,683 long-reads with 6.6 GB bases (Supplementary Figure 13). We aligned the data against a reference genome (hg38) using minimap2[35], and 63% long-reads (73% bases) are mapped. Then, we analyzed this ONT RNA-seq data with LIQA for genes with at least two isoforms.

We considered two ways to benchmark the performance of LIQA. First, we used PennSeq to analyze short-read sequencing data for the same AML sample and treated the estimates as gold standard. This dataset included 440M short read with 150bp in length. Figure 5(A) shows the scatter plots of isoform relative abundance estimates between long- and short-reads data. Spearman’s correlation coefficients were calculated. We found that correlation was improved significantly for genes with at least 50 reads compared to all genes without filtration. Then, we examined the major isoforms (with the highest expression level in a gene) inferred by LIQA. As shown in Figure 5(B), long-read and short-read shared consistent estimates for the major isoforms. This is not surprising because major isoforms were more likely to be sequenced, leading to higher read coverage at unique exonic regions. Second, we visually examined the read coverage plots at unique exonic regions with at least 100 reads to benchmark the performance of LIQA. We generated sashimi plots for two randomly selected genes, *EOGT* and *RRBP1* (Figure 5(C)). For gene *EOGT*, the read coverage ratio between exons in red and green squares suggests that isoforms NM_103826 and NM_001278689 expressed much higher than NM_173654. This is consistent with LIQA’ estimates, with relative abundance of NM_173654 less than 0.01. A similar pattern is observed for gene *RRBP1*, where isoform NM_004587 (relative abundance estimates = 0.68) is the major isoform. Results from this AML data demonstrate the robust performance of LIQA to 3’ coverage biases.

**Figure 5.**
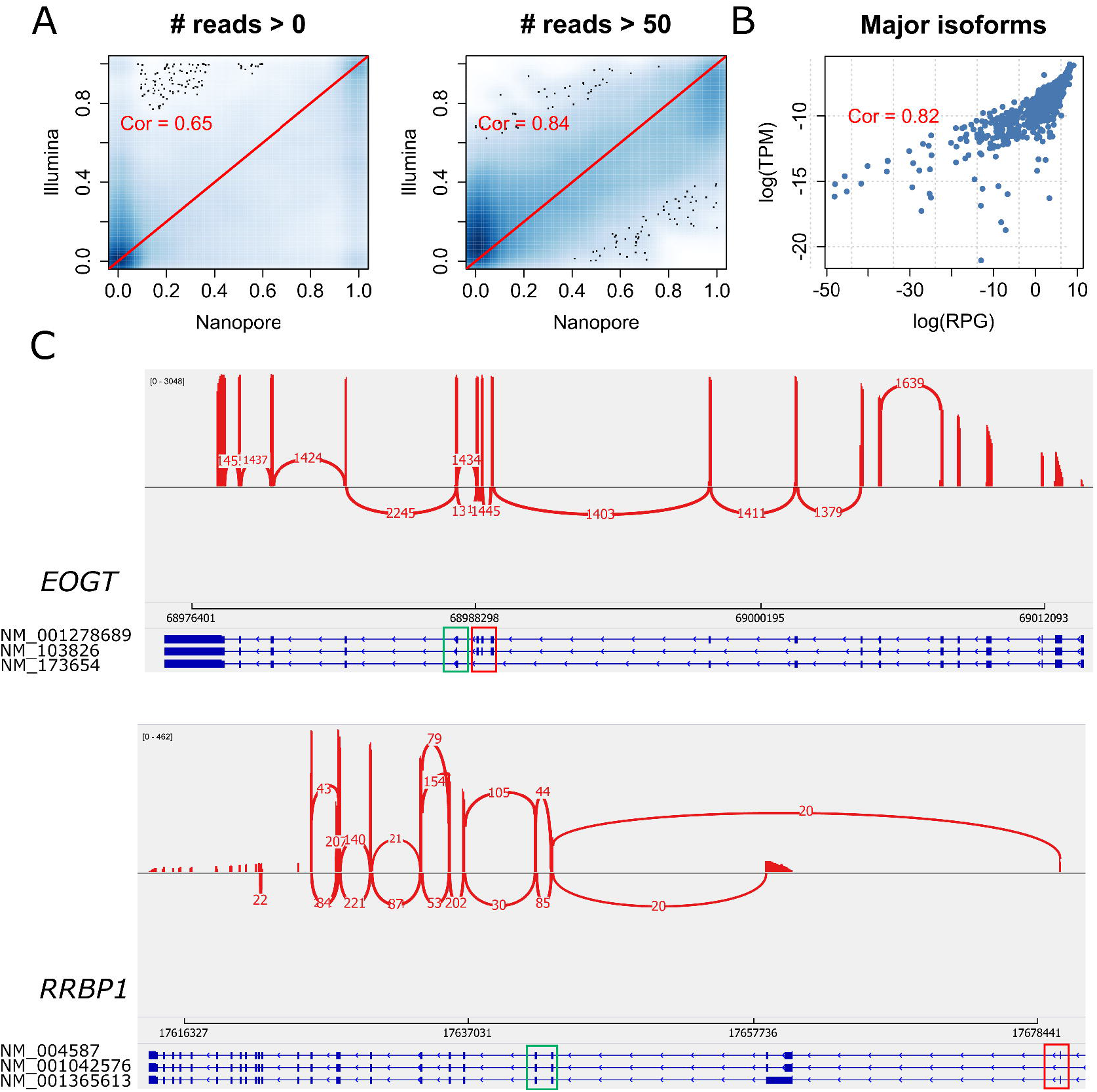
Performance of LIQA using AML data. (A) Scatter plots of estimated isoform relative abundances using long-read data (LIQA) versus short-read data (PennSeq) for all genes (left) and genes with at least 50 read coverage (right). (B) Scatter plot of estimated isoform-specific expression using long-read data (LIQA) versus short-read data (PennSeq) in log-scale for all major isoforms. (C) Examination of isoform usage inferred by LIQA. Sashimi plots of gene *EOGT* and *RRBP1*. Informative exonic regions were in green and red squares.

### Application to PacBio data on esophageal squamous epithelial cell (ESCC)

Next, we evaluated the performance of LIQA in differential alternative splicing (DAS) detection using an RNA-seq dataset generated from esophageal squamous epithelial cell (ESCC)[40]. This dataset includes PacBio SMRT reads generated from normal immortalized and cancerous esophageal squamous epithelial cell lines. The RNA-seq data were downloaded from Gene Expression Omnibus (PRJNA515570). We applied LIQA to detect differential isoform usage between normal-like and cancer cells. Known splicing differences in existing studies were treated as ground truth to evaluate LIQA’s performance in characterizing isoform usage across samples. In addition, short-read data from these two samples were sequenced using the Illumina platform, which allows us to compare the consistency and accuracy of DAS detection between long-read and short-read data.

Employing LIQA and PennDiff[41], PacBio and Illumina data were analyzed to detect DAS events, which are classified into different types, such as skipped exon (SE), alternative 5’ splice site (A5SS), alternative 3’ splice site (A3SS), mutually exclusive exon (MXE) and retained intron(RI). Our results showed that SE is the most frequent type of event among detected DAS between normal-like and cancerous cells, followed by RI, A5SS and A3SS. MXE is the most infrequent splicing type. As shown in Figure 6(A), detected DAS events by long- and short-read share strong association at both exon and gene level (Cramer’s V > 0.5). Also, the concordance rate between long and short read data is greater than 98%. Compared to short-read, long-read data shows preference in detecting more differential splicing events at both exon and gene level. This is not surprising because read coverage heterogeneity, which might bias DAS detection, is alleviated in long-read data by capturing full-length transcript in each read.

**Figure 6.**
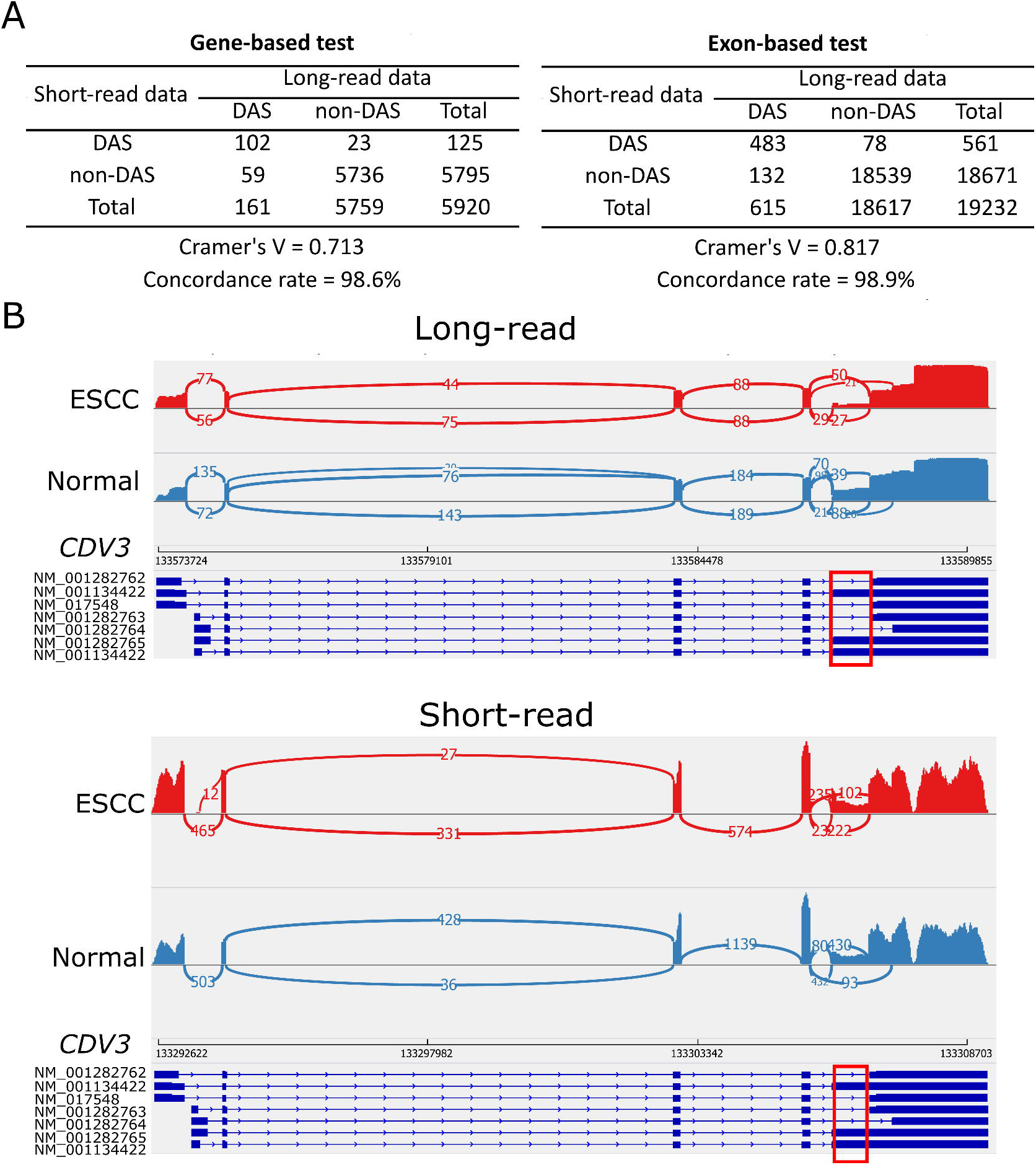
Performance of LIQA using ESCC data. (A) DAS detections between long- and short-read data. Consistency of detected DAS events between long- and short-read data were quantified using Cramer’s V and concordance rate. (B) Examination of AS exon usage inferred by LIQA (long-read) but missed by PennDiff (short-read). Sashimi plots of gene *CDV3*. Informative exonic regions were red squares.

The expression of alternatively spliced isoforms from gene *CDV3* shows difference in cancerous ESCC cells compared to non-cancerous[42, 43]. Figure 6(B) provides the sashimi plots of a DAS exon at gene *CDV3* detected by LIQA, but was missed by PennDiff using short-read data. From long-read data, it is clear that the relative expression of exon in the red square is lower in cancerous cells than normal-like ESCC. However, this event is missed by short-read data. The read coverage difference between normal-like and cancerous ESCC in sashimi plots indicates the less usage of isoforms (NM_001134422, NM_0011134423, NM_001282765) which include this exon in ESCC cells, suggesting better performance of long-read data.

## Discussion

Accurate estimation of isoform-specific gene expression is a critical step for transcriptome profiling. The emergence of long-read RNA-seq has made it possible to discover complex novel isoforms and quantify isoform usage based on full-length sequenced fragments without amplification bias. However, there are still issues for long-read data, which if not taken into account, can affect the estimations. The major challenges in the analysis of long-read RNA-seq data are the presence of high error rate and potential coverage bias. In this article, we propose LIQA, a statistical method that allows read-specific weight in estimating isoform-specific gene expression. The central idea of our method is to extract error rate information and model non-uniformity read coverage distribution of long-read data. LIQA is the first long-read transcriptomic tool that takes these limitations of long-read RNA-seq data into account. Results of our simulation study and analyses of real data demonstrated that LIQA is more effective in bias correction than the limited existing approaches (Supplementary Table 2, 3).

However, we note that there is still room to improve LIQA. LIQA is computationally intensive because the approximation of non-parametric Kaplan-Meier estimator of function *f*(*L*_*r*_)relies on empirical read length distribution and the parameters are estimated using EM-algorithm. Based on the analysis of the UHR and AML data, we found that running LIQA is slower than FLAIR and Mandalorion (Supplementary Table 1). Currently, we are evaluating the impact of possible parametric functions such as exponential or Weibull distributions for read distribution modeling. This will sacrifice the robustness of isoform expression estimates but the running time can be significantly reduced. We believe it may be worth making this trade-off between computational efficiency and estimation accuracy for LIQA.

We have benchmarked the performance of LIQA with the use of minimap2 for long-read alignment, while there have been several approaches supporting RNA-seq long-read alignment, such as STAR[37], GMAP[44], BLAT[45], BBMap (https://sourceforge.net/projects/bbmap/), and GraphMap2[46]. LIQA can take SAM or BAM files generated from any listed aligner as input. Nevertheless, we recognize that it is important to evaluate whether LIQA’s superior performance is robust to different aligners. Therefore, we plan to explore more long-read aligner options and settings to benchmark LIQA in the future.

As LIQA is EM-algorithm-based, the robustness to parameter initialization is a potential issue. Read-specific weight of LIQA extracts more information from observed data than direct read-count strategy as implemented in Mandalorion and FLAIR. Especially, more read coverage is needed for stable approximation of function *f*(*L*_*r*_). For genes with limited reads coverage (less than 5), the likelihood function of LIQA will be flattened, then optimal points are harder to be reached by EM-algorithm and estimates may be sensitive to initial values of parameter. Therefore, the sensitivity of LIQA to parameter initialization should be further evaluated and improved.

With full-length transcript sequencing, long-read RNA-seq data (ONT and PacBio) are expected to facilitate transcriptomic studies by offering number of advantages over short reads. For PacBio, HiFi reads are generated with circular consensus sequencing (CCS) using single-molecule consensus, which increases their accuracy over traditional multi-molecule consensus. Compared to Nanopore sequencing, this protocol yields much lower per-base error rate compared to Nanopore sequencing, but potentially shorter reads. Smaller read length may introduce much larger biases in 5’ or 3’ coverage ratio, which requires further adjustment for LIQA to derive more accurate isoform expression estimates. LIQA has custom settings that allow users to flexibly adjust such parameters to handle these platforms. Compared to PacBio (either with traditional library or HiFi library preparation protocols), ONT may be a more promising platform in quantifying isoform expression while generating data with much higher error rate. This is because ONT is currently more affordable with lower per-based cost of data generation, and sequencing data with high read coverage can improve estimation accuracy of isoform usage. For ONT RNA-seq, there are two types: direct mRNA sequencing and cDNA sequencing. Compared to direct mRNA sequencing, cDNA sequencing allows samples to be amplified and requires less amounts of starting materials. Our studies showed that the decrease of read coverage had less impact on LIQA compared to other existing approaches.

In summary, long-read RNA-seq data offer advantages and can help us better understand transcriptomic variations than short-read data. However, better utilizing informative single molecule sequencing read is not straightforward. LIQA is a robust and effective computational tool to estimate isoform-specific gene expression from long-read RNA-seq data. With the increasing adoption of long-read RNA-seq in biomedical research, we believe LIQA will be well-suited for various transcriptomics studies and offer additional insights beyond gene expression analysis in the future.

## Methods and materials

### Complete likelihood function of LIQA

Given a gene of interest, let ***R*** denote the set of reads that are mapped to the gene of interest, and ***I*** denote the set of known isoforms. For a specific isoform *i* ∈ ***I***, let *θ*_*i*_ denote its relative abundance, with 0 ≤ *θ*_*i*_ ≤ 1 and ∑_*i*∈***I***_ *θ*_*i*_ =1 and *l*_*i*_ denote its length. For each single-molecule long-read *r*, let *L*_*r*_ denote its length. The probability that a read originates from isoform *i* is *P* (iso.= *i*) *= θ*_*i*_. For read-isoform assignment, LIQA accounts for incorrect alignment at splice site. We define parameter ***Z***_***R***,***I***_ as a |*R*| *×* |*I*| a read-isoform compatibility matrix with ***Z***_***R***,***I***_ (*r,i*) = 1 if long-read *r* is generated from a molecule that is originated from isoform *i* (number of mismatch base pairs < 20 bp instead of exact match), and ***Z***_***R***,***I***_ (*r,i*) = 0 otherwise. For isoform quantification, our goal is to estimate Θ = {*θ*_*i*_,*i* ∈ ***I***} based on RNA-seq long-reads mapped to the gene.

With the notation above, the complete data likelihood of the RNA-seq data can be written as

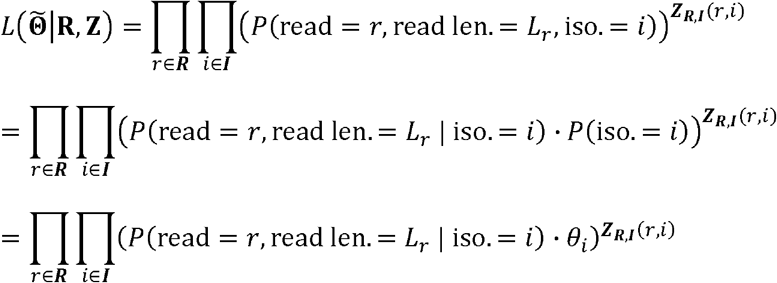

This formula is based the fact that given the isoform origin, the probability of observing read alignment can be inferred. The conditional probability of read *r* derived from isoform *i* with length *L*_*r*_ is

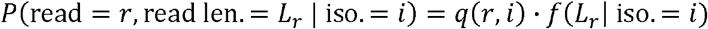

where *q*(*r,i*) is isoform-specific read quality score and *f*(*L*_*r*_ | iso. = *i*) is isoform-specific read length probability. Essentially, we quantify isoform relative abundance with weighted read assignment. To account for the error-prone manner of Nanopore sequencing data, we consider isoform-specific read quality score 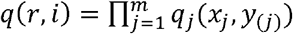 where *x* is the sequence of the long-read *r,y* is the sequence of the corresponding isoform *i* in the reference genome, and *q*_*j*_ (*a,b*) is the probability that we observe base *a* at position *j* of the read given that the true base is *b*, which can be calculated as 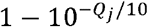, with *Q*_*j*_ being the per-based Phred quality score at position *j*.

### Estimation of isoform-specific read length probability *f*(*L*_*r*_ | iso. = *i*)

Because read length *L*_*r*_ is not fixed and short prone in Nanopore sequencing, we treat it as a random variable with right skewed distribution density function *f* (·). Given an isoform, existing long-read methods assume fixed read length for all sequenced read, and this is equivalent to setting *f*(*L*_*r*_) at 1. However, this assumption does not hold as recent studies suggest that potential 3’ coverage bias exists in long-read RNA-seq data [24, 32, 47]. To offer flexibility in modeling read length distribution, we employ a nonparametric approach. For all long-reads mapped to the genome, we categorize them into two groups: complete reads and truncated reads. Accounting for misalignment due to high error rate, the read is treated as complete when the distance between its ending alignment position and any known isoform 5’ end is less than a tolerance threshold (default = 20 bp) (Figure 1(A)). This indicates that this read is completely sequenced from a known isoform. Otherwise, the read is considered as truncated. The presence of truncated reads is due to incomplete sequencing or novel isoforms. As known annotated isoforms are treated as gold standard during estimation, we assume true length of truncated read is censored. Given the observed lengths of all complete and truncated reads, we fit them into a survival model, a natural modeling approach for censored data (Supplementary Figure 9, 10, 11). Function 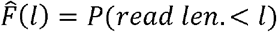 can be estimated based on Kaplan-Meier estimator[48], hence we have 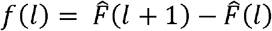.

Given a gene of interest with ***I*** = {isoform *i*: 1 ≤ *i* ≤ *I*}, isoform-specific read length probability *f*(*L*_*r*_ | iso. = *i*) can be written as

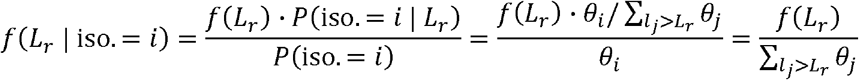

This isoform-specific read length probability *f*(*L*_*r*_ | iso. = *i*) captures the sequencing biases due to fragmentation during library preparation or pore-blocking for nanopore data.

### Quantification of isoform expression level

Given that isoform indicators ***Z***_***R***,***I***_ (*r,i*) for some reads are not observed from read data, **Θ** are estimated using EM algorithm. Then, we have isoform relative abundance 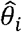. In addition to relative abundance, it is also important to quantify the absolute expression level of an isoform. At gene level, we consider read per gene per 10K reads (RPG 10K) as the standard for long-read RNA-seq data. RPG is defined as RPG = *N*/10^4^ where *N* is the number of reads mapped to the gene of interest. With this concept, we estimate the expression level of a particular isoform by replacing *N* with estimated number of long-reads originated from isoform 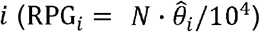.

### Parameter estimation using the EM algorithm

The complete data likelihood is

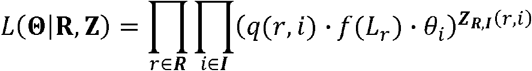

and the update procedure of the EM algorithm is as follows:

**E-step:** We calculate function

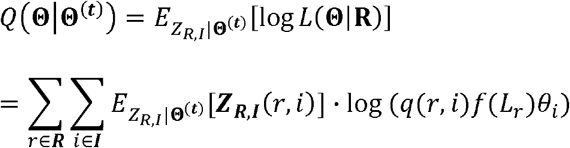

where 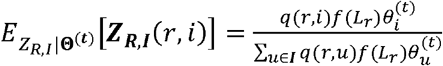.

**M-step:** We maximize function *Q*(**Θ**|**Θ**^(**t**)^)and have

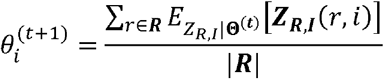

The EM algorithm consists of alternating between the E- and M-steps until convergence. We start the algorithm with **Θ**^(**0**)^ assuming all isoforms are equally expressed and stop when the log likelihood is no longer increasing significantly.

### Detection of differential alternative splicing (DAS) with LIQA

The relative abundance of an isoform takes values between 0 and 1. Therefore, we assume it follows a beta distribution, which is well known as a flexible distribution in modeling proportion because its density can have different shapes depending on the values of the two parameters that characterize the distribution. i.e. *θ*_*i*_ ∼Beta (*μ*_*i*,_ *ϕ*_*i*_). The expected value and variance of *θ*_*i*_ are

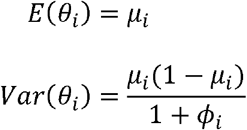

To detect splicing difference of isoform *i* between two groups of samples, we utilized beta regression model with *ϕ*_*i*_ as precision parameter. We apply logit link function and have the model

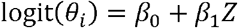

where *Z* is the condition indicator (1 for case; 0 for control), *β*_0_ and *β*_1_ are coefficient parameters.

Since the isoform relative abundances of isoforms within the same gene are correlated, a robust and flexible model is needed when comparing them between conditions at gene level. To account for this, we utilize Gaussian copula regression model to test splicing difference significance between conditions of correlated isoform relative abundances. The separation of marginal distributions and correlation structure makes Gaussian copula regression versatile in modeling non-normal dependent observations. Therefore, the joint distribution of isoform relative abundances from the same gene is given by

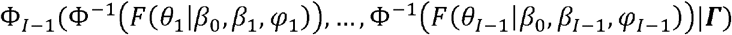

where *φ*_*i*_ is the dispersion parameter of the marginal generalized linear model for isoform *i*. *ϕ*_*I*−1_(|***Γ***) is the cumulative distribution function of multivariate normal random variables with *I* – 1 dimensions and correlation matrix ***Γ***. We choose to use exchangeable correlation structure for ***Γ***. Given regression models above, we can detect DAS both for at isoform level and at gene level. For isoform *i*, we test *H*_0_: *β*_1_ = 0 vs *H*_1_: *β*_1_ ≠ 0 to determine splicing change between conditions. For gene *g*, we test 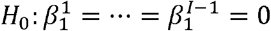 vs 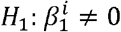 for any 1 ≤ *i* ≤ *I –* 1.

### Nanopore direct mRNA sequencing of universal human reference RNA-seq data

Universal human reference (UHR) RNA comprises of mixed RNA molecules by a diverse set of 10 cancer cell lines with equal quantities of DNase-treated RNA from adenocarcinoma in mammary gland, hepatoblastoma in liver, adenocarcinoma in cervix, embryonal carcinoma in testis, glioblastoma in brain, melanoma, liposarcoma, histocytic lymphoma in histocyte macrophage, lymphoblastic leukemia and plasmacytoma in B lymphocyte. This reference sample from MicroArray Quality Control (MAQC)[49-51] project has been utilized in many studies. For example, Gao et al[52] sequenced this UHR RNA sample and treated it as reference to measure the technical variations of scRNA-seq data. Also, the qRT-PCR measurements of gene or isoform expressions from this sample were used to benchmark and optimize computational tools[17, 53-56]. In this study, we used GridION Nanopore technique to sequence mRNA directly, and used Guppy for base calling. In total, we generated 476,000 long-reads with 557 MB bases. We aligned the UHR RNA-seq data against a reference genome (hg38) using minimap2[35], and 95% long-reads (89% of total bases) are mapped, demonstrating very high sequencing and basecalling quality. qRT-PCR measurements were downloaded and treated as ground truth to compare the performance between LIQA, FLAIR, Mandalorion, CEM, Cufflinks and RD.

### Chi-squared goodness of fit statistics of read coverage uniformity

Given an isoform of interest, let *l* denote the length and *O*_*i*_ denote observed read coverage count at base pair position *i*. Total sequencing depth of this isoform *S* = ∑_1≤*i*≤*l*_ *O*_*i*._ Under the uniform read coverage assumption, the expected read coverage count at each base pair position *E*_*i*_ = *S*/*l*. We apply Chi-squared goodness of fit statistics to measure the difference between observed read coverage and uniform read distribution. The test statistics is

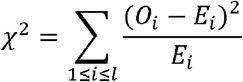

The higher value of *χ*^2^ indicates that observed read coverage deviates more from uniform read distribution. We calculated *χ*^2^ for each isoform, then divided them into two categories based on median of the corresponding measure (less uniform: *χ*^2^> median, more uniform *χ*^2^ < median) to evaluate the impact of read coverage distribution on isoform expression quantification.

### Statistical test to compare performance of different methods

We simulated ONT RNA-seq data 20 times to assess the statistical significance when comparing the performance of different methods. Each dataset includes 5 million (5M) reads. We also down-sampled 3 million (3M), 1 million (1M) and 0.5 million (0.5M) reads for the simulated data to evaluate the impact of sequencing depth on performance improvement of LIQA. We ran all methods with the same set of simulated aligned data in BAM format as input and calculated the Spearman’s correlation of TPM and relative abundance between true and estimated values. Based this metric from 20 simulated datasets, we conducted pairwise comparison of performance difference between all methods using paired Z-test. Mean difference, standard deviations, test statistics and P-values were calculated. Moreover, we conducted likelihood ratio test to compare different models of LIQA (full model, read length model, read quality model). Likelihood ratio test statistic Q = −2(log*L*_*B*_ − log*L*_*A*_) where *L* is the optimized likelihood function based on different models.

## Data availability

The direct mRNA sequencing data on UHR has been deposited and available at BioProject database (PRJNA639366). The cDNA sequencing data on a patient with cancer has been deposited and available at BioProject database (PRJNA640456). The simulation data used in our study can be reproduced using code provided in the LIQA software repository and NanoSim version 2.0.0.

## Acknowledgements

We thank the Wang lab members for insightful comments and for testing the software tools. We also thank the developers of the NanoSim software tool, and the generators of the short-read and qRT-PCR results on the UHR datasets for making the data publicly available for benchmarking studies. We thank three anonymous reviewers for their constructive comments and suggestions on benchmarking studies.

## Funding

This study is supported by NIH/NIGMS grant GM132713 and the CHOP Research Institute.

## Notes

### Competing Interest Statement

The authors have declared no competing interest.

