## Supplemental material for "LIQA: Long-read Isoform Quantification and Analysis"

**Supplementary Figures**

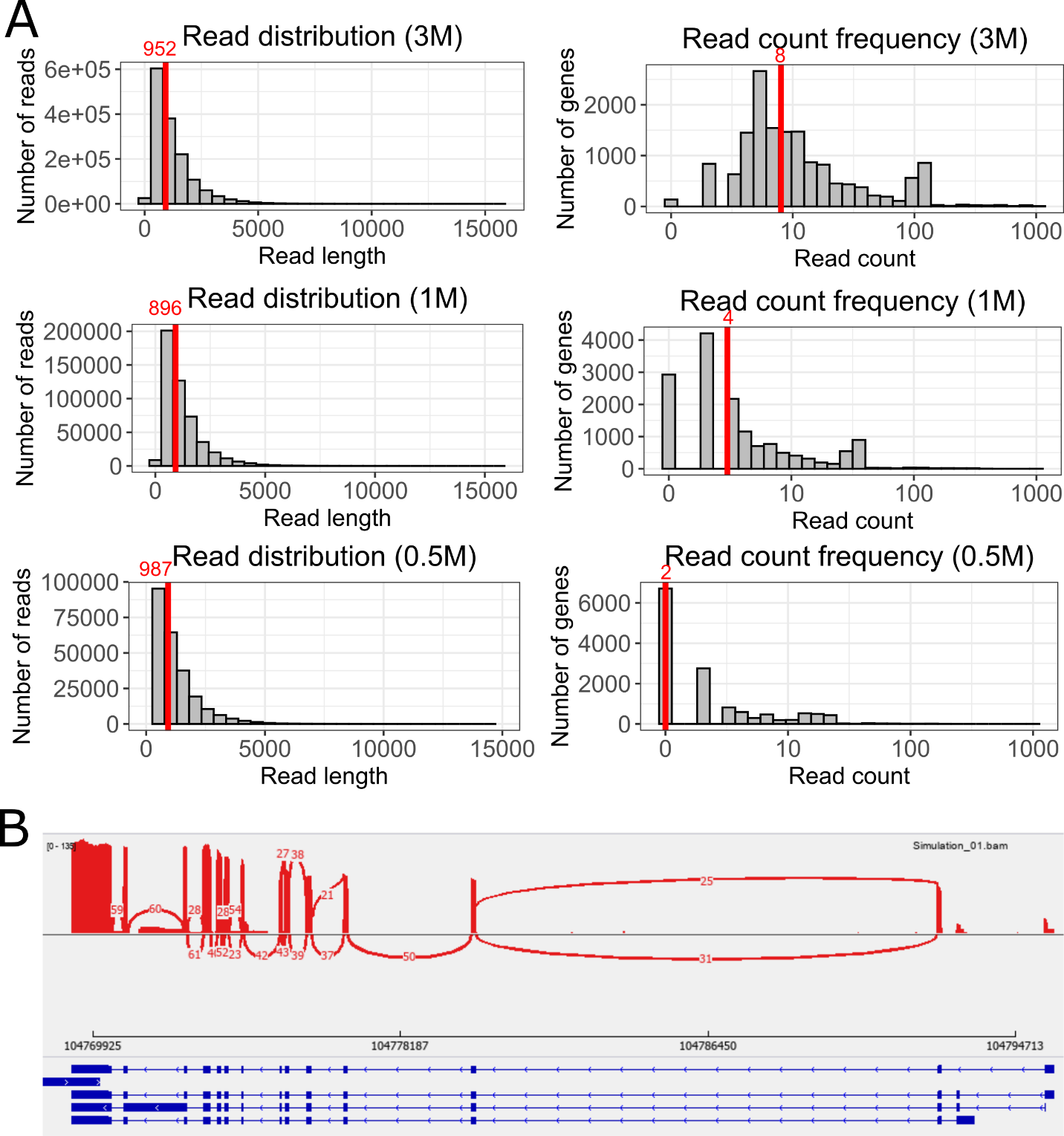

**Supplementary Figure 1. Simulated data summary.** (A) Characteristics of simulated data with different depths (3M, 1M, 0,5M). Read length distribution (left panel) and read count distribution by genes in log-scale (right panel). (B) Sashimi plot of simulated long-read RNA-seq data using NanoSim.

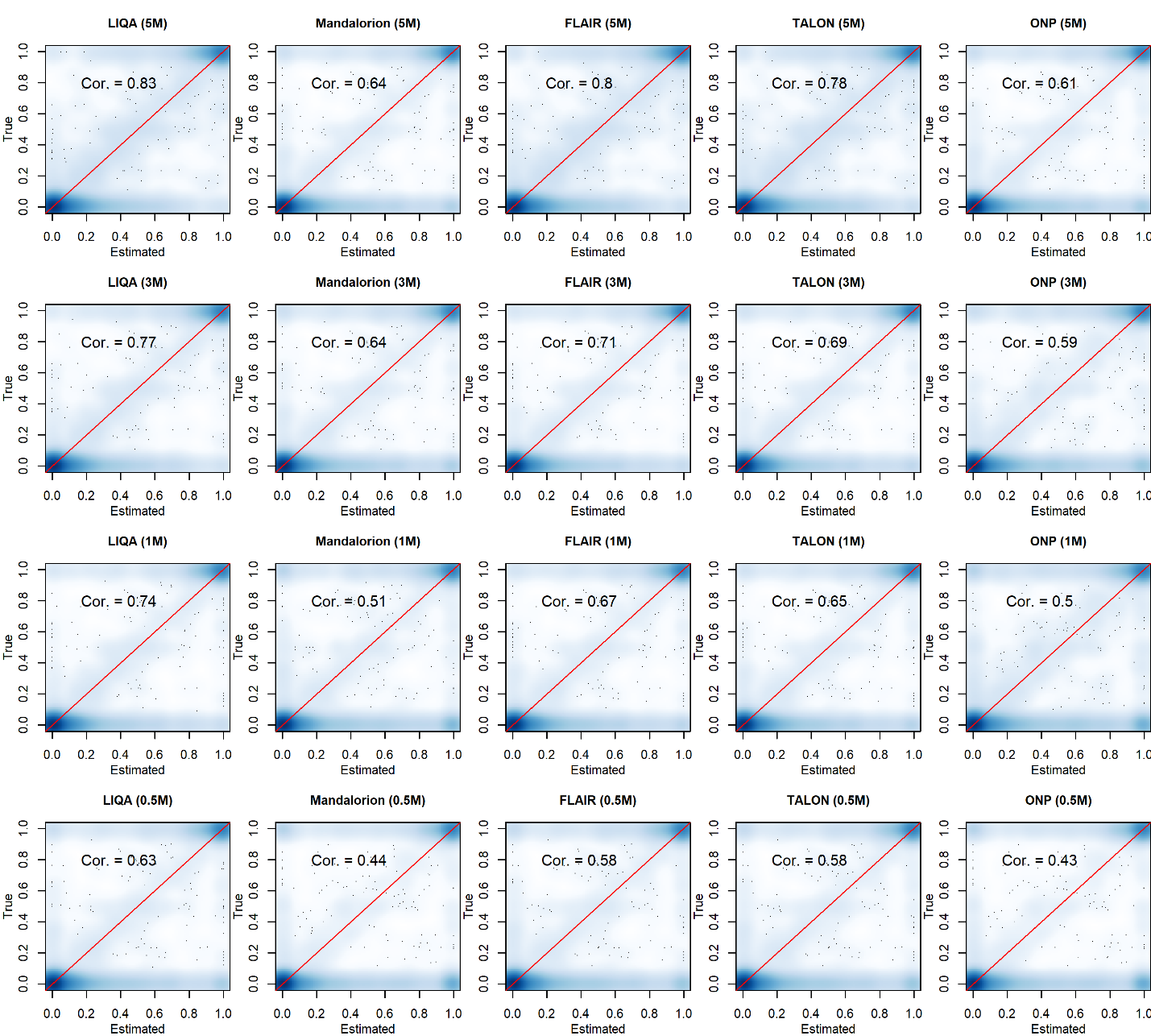

**Supplementary Figure 2. Performance comparison between methods based on simulated data.** Scatter plots of estimated isoform relative abundance vs true relative abundance for the simulated data at different read depths (5M, 3M, 1M, 0.5M).

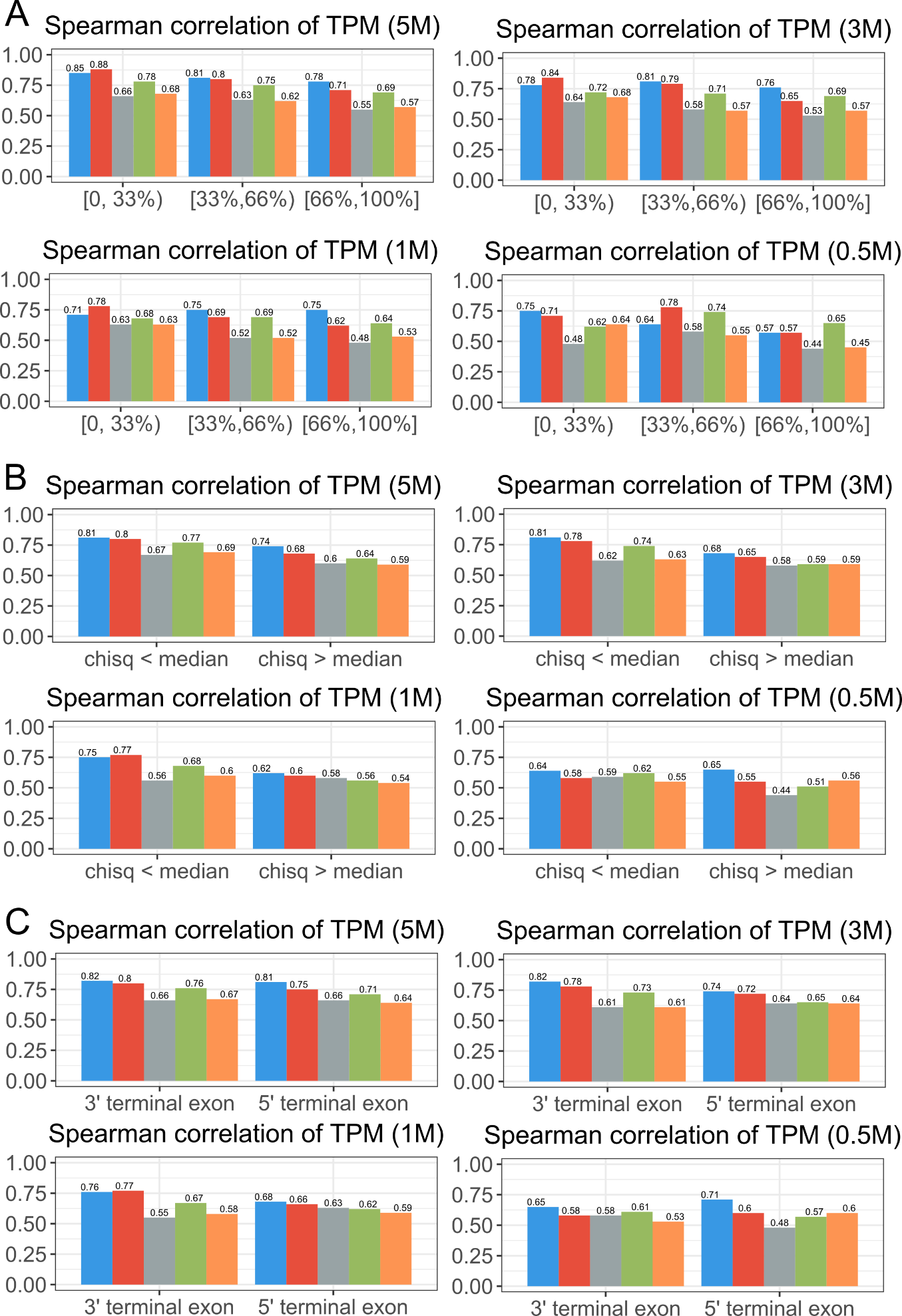

**Supplementary Figure 3. Evaluation of robustness to read length bias and low read quality at different read coverage levels using LIQA (blue), FLAIR (red), Mandalorion (gray), TALON (green) and ONP (yellow).** (A) Simulation results spearman correlations between estimated and true TPM for isoforms with different lengths (length < 33% quantile, 33% ≤ length < 66% quantile, length ≥ 66%). (B) Spearman correlations between estimated and true TPM. Isoforms are stratified by the chi-squared goodness of fit statistic for uniformity. (C) Spearman correlations between estimated and true TPM for 3’ and 5’ terminal exons. Summary statistics were calculated for each method at different read depths (5M, 3M, 1M, 0.5M) to compare the robustness to read coverage bias.

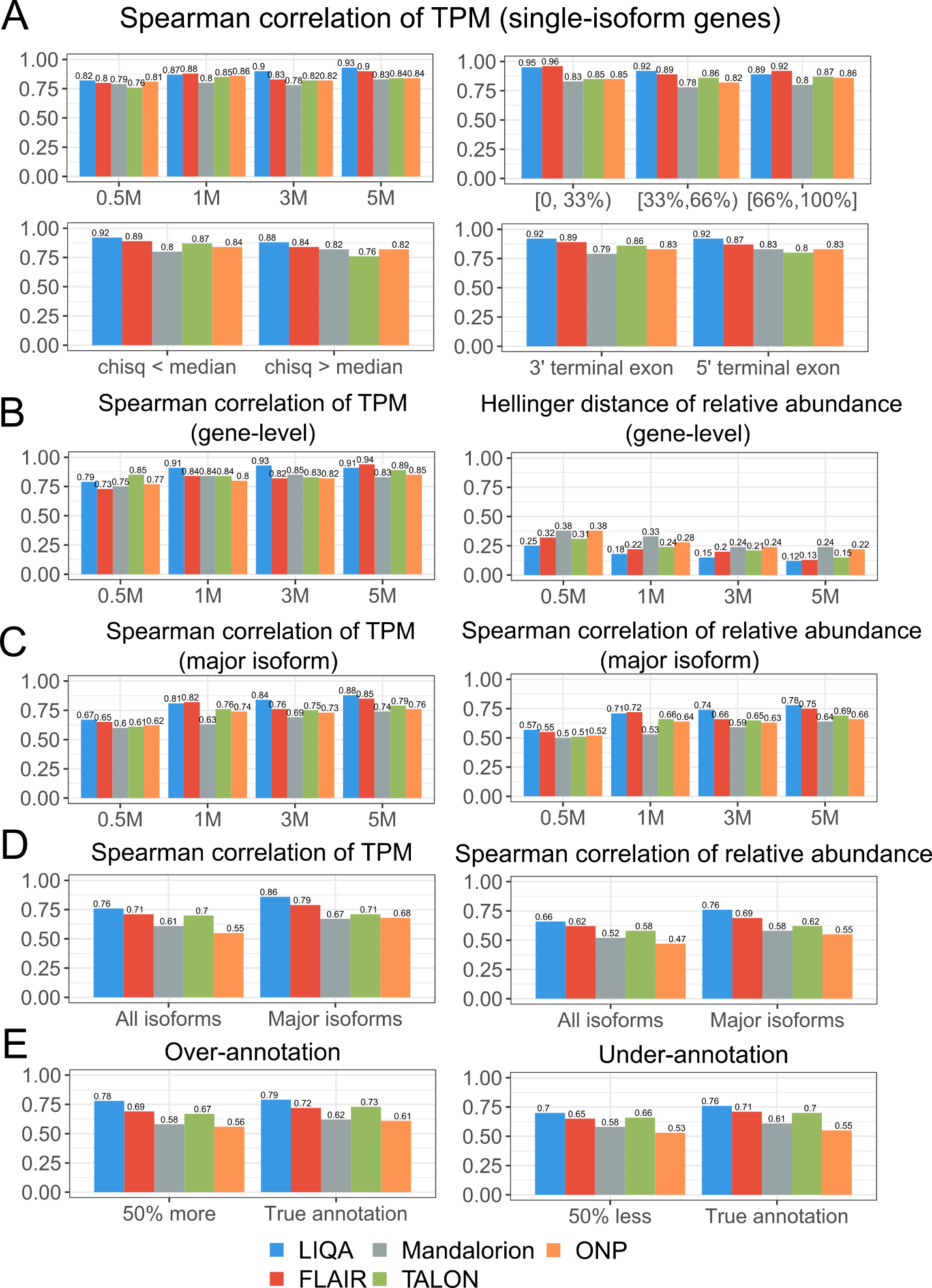

**Supplementary Figure 4. Simulation study of gene-level quantification and robustness to isoform misannotation using LIQA (blue), FLAIR (red), Mandalorion (gray), TALON (green) and ONP (yellow).** (A) Estimation accuracy comparison based on single-isoform genes using spearman correlations between estimated and true TPM at different read coverage (top-left), for isoforms with different lengths (length < 33% quantile, 33% ≤ length < 66% quantile, length ≥ 66%) (top-right), for isoforms are stratified by the chi-squared goodness of fit statistic for uniformity (bottom-left) and for 3’ and 5’ terminal exons (bottom-right). (B) Spearman correlations between estimated and true gene-level TPM at different read coverage (left). Average Hellinger distance between estimated and true relative abundance of isoforms at gene-level (right). (C) Spearman correlations between estimated and true gene-level TPM (left) and relative (right) abundance for major isoforms. (D) Spearman correlations between estimated and true gene-level TPM (left) and relative (right) abundance for all isoforms using GENCODE v37. (E) Evaluation of robustness to isoform over- (left) and under-annotation (right) using GENCODE v37.

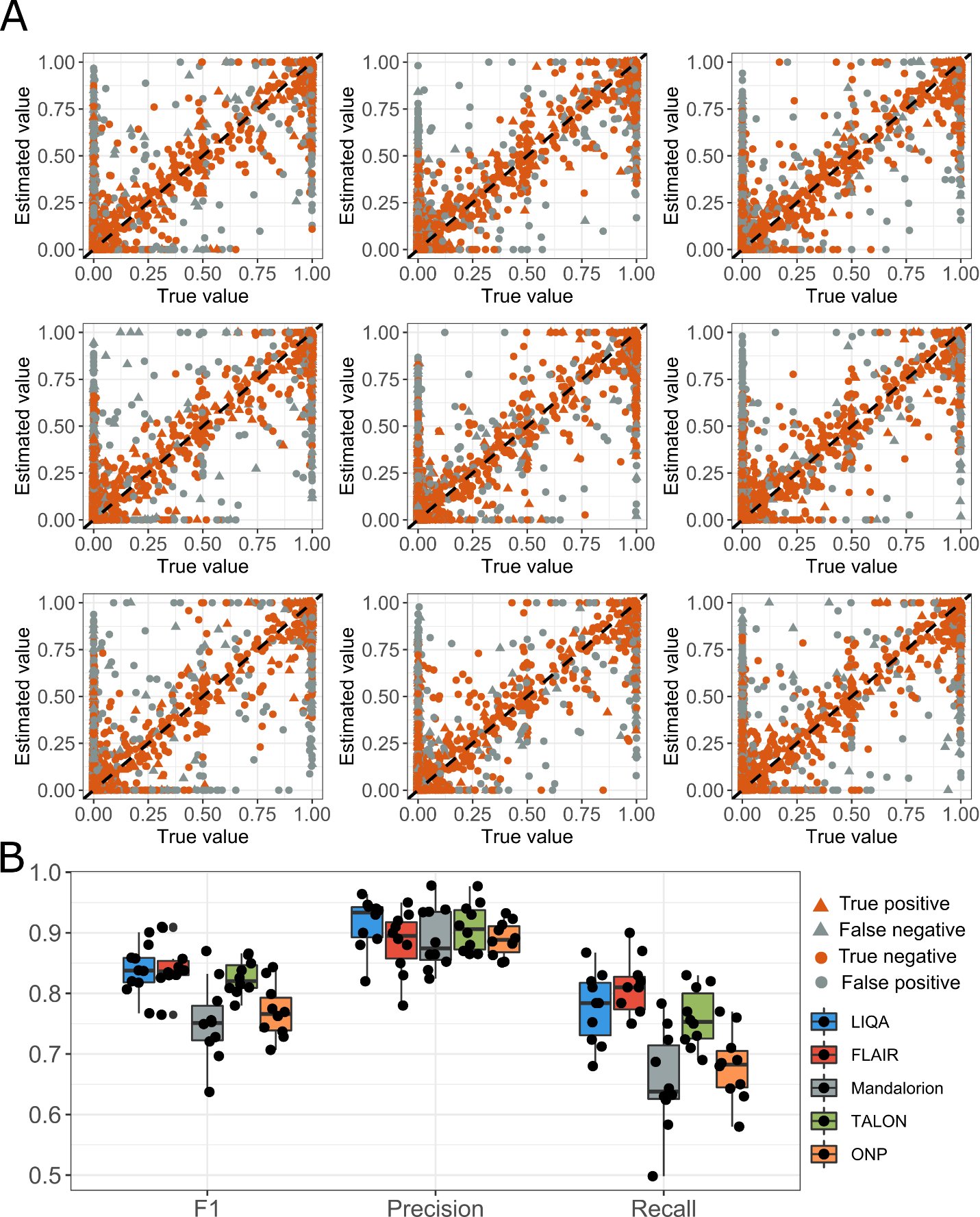

**Supplementary Figure 5.** **Differential splicing analysis results based on simulated data.** (A) Scatter plots of true and estimated isoform relative abundance of LIQA based on 9 DAS simulation datasets. LIQA’s prediction and ground truth are marked for each isoform. (B) Summary statistics (recall, precision and F1 score) of DAS event detection analysis for 10 simulations

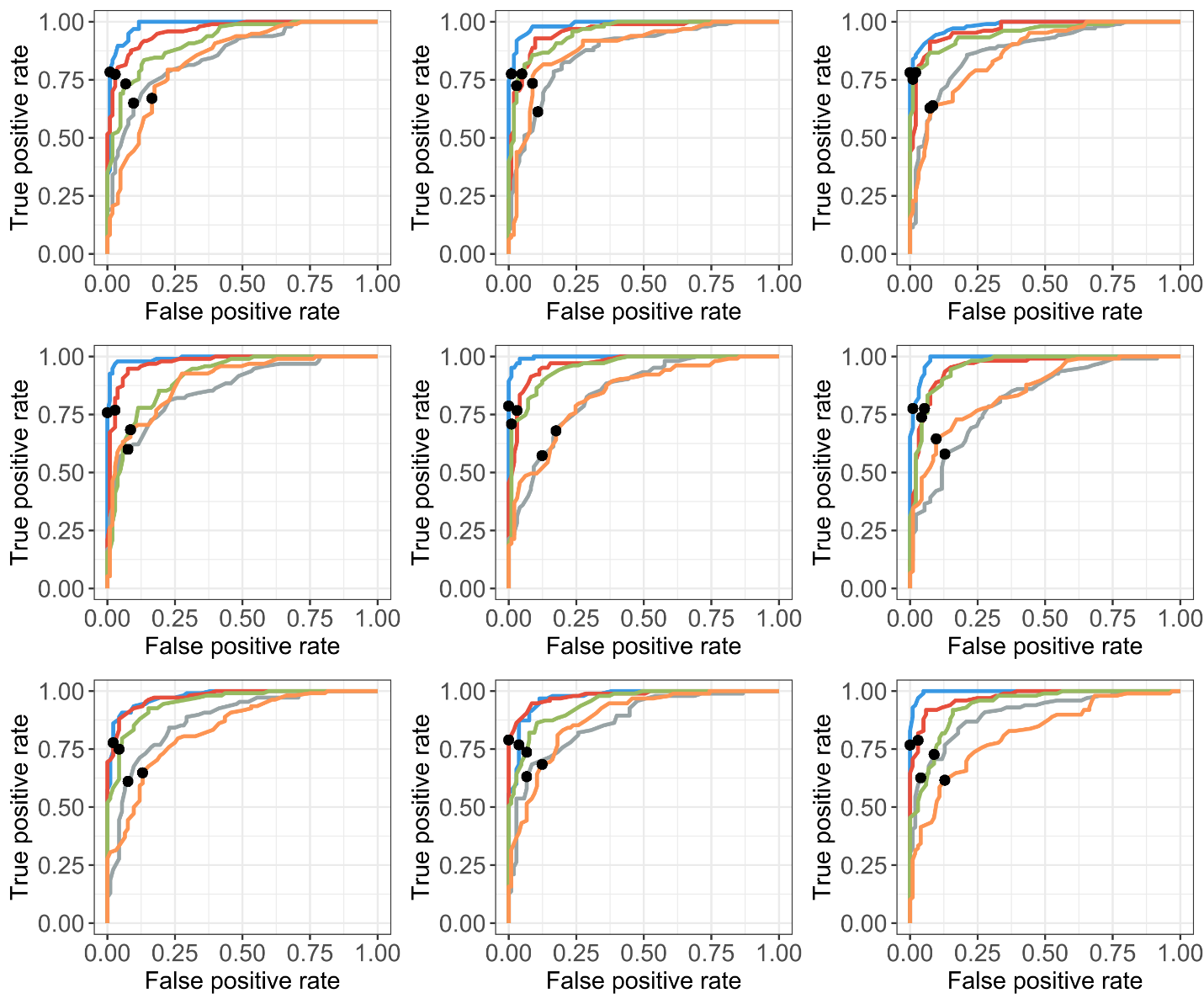

**Supplementary Figure 6.** **Differential splicing analysis results based on simulated data using LIQA (blue), FLAIR (red), Mandalorion (gray), TALON (green) and ONP (yellow).** ROC curve of different methods in detection DAS genes based on 9 simulation datasets. Threshold FDR<0.05 is highlighted using black dot.

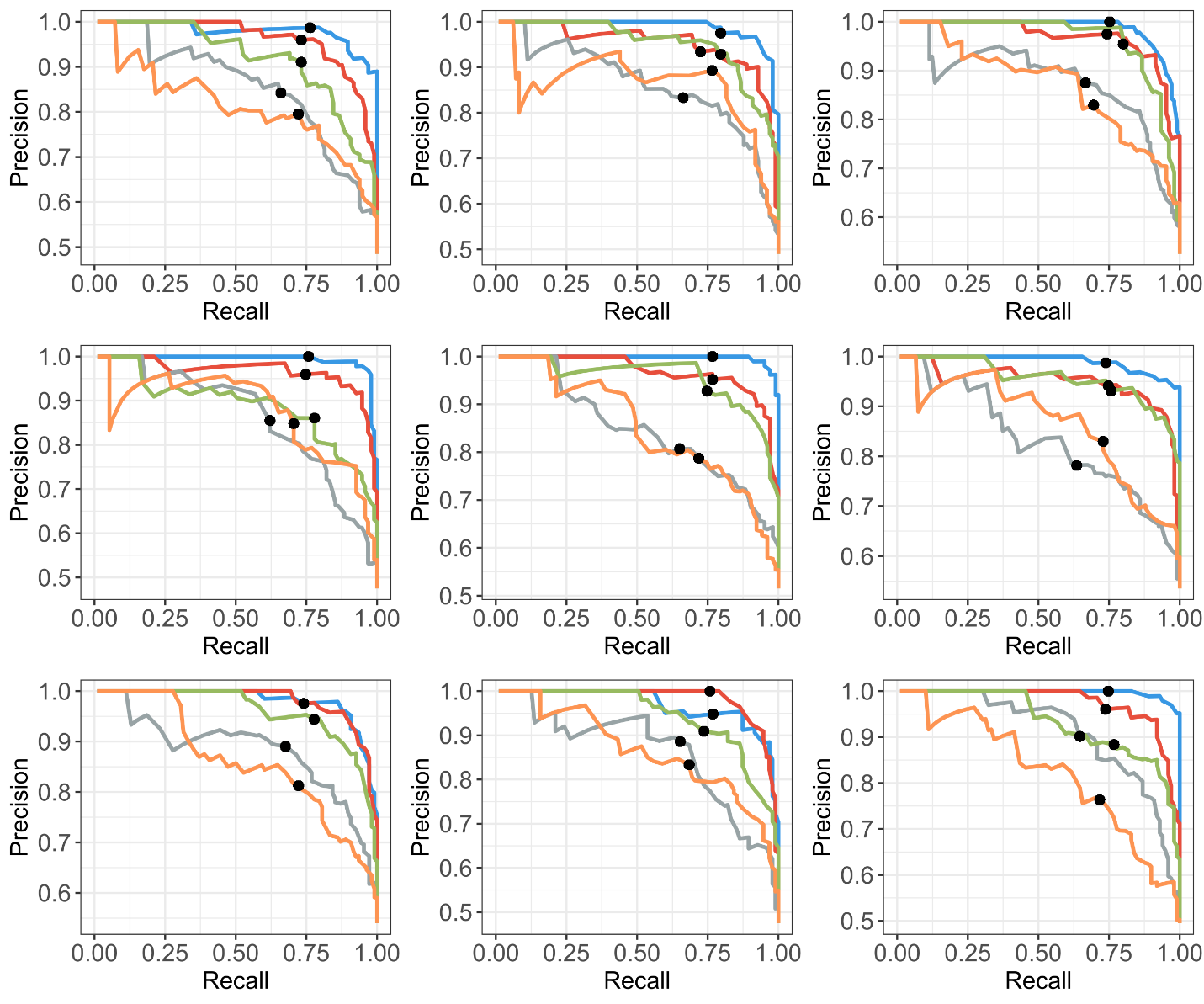

**Supplementary Figure 7.** **Differential splicing analysis results based on simulated data using LIQA (blue), FLAIR (red), Mandalorion (gray), TALON (green) and ONP (yellow).** Precision-recall curve of different methods in detection DAS genes based on 9 simulation datasets. Threshold FDR<0.05 is highlighted using black dot.

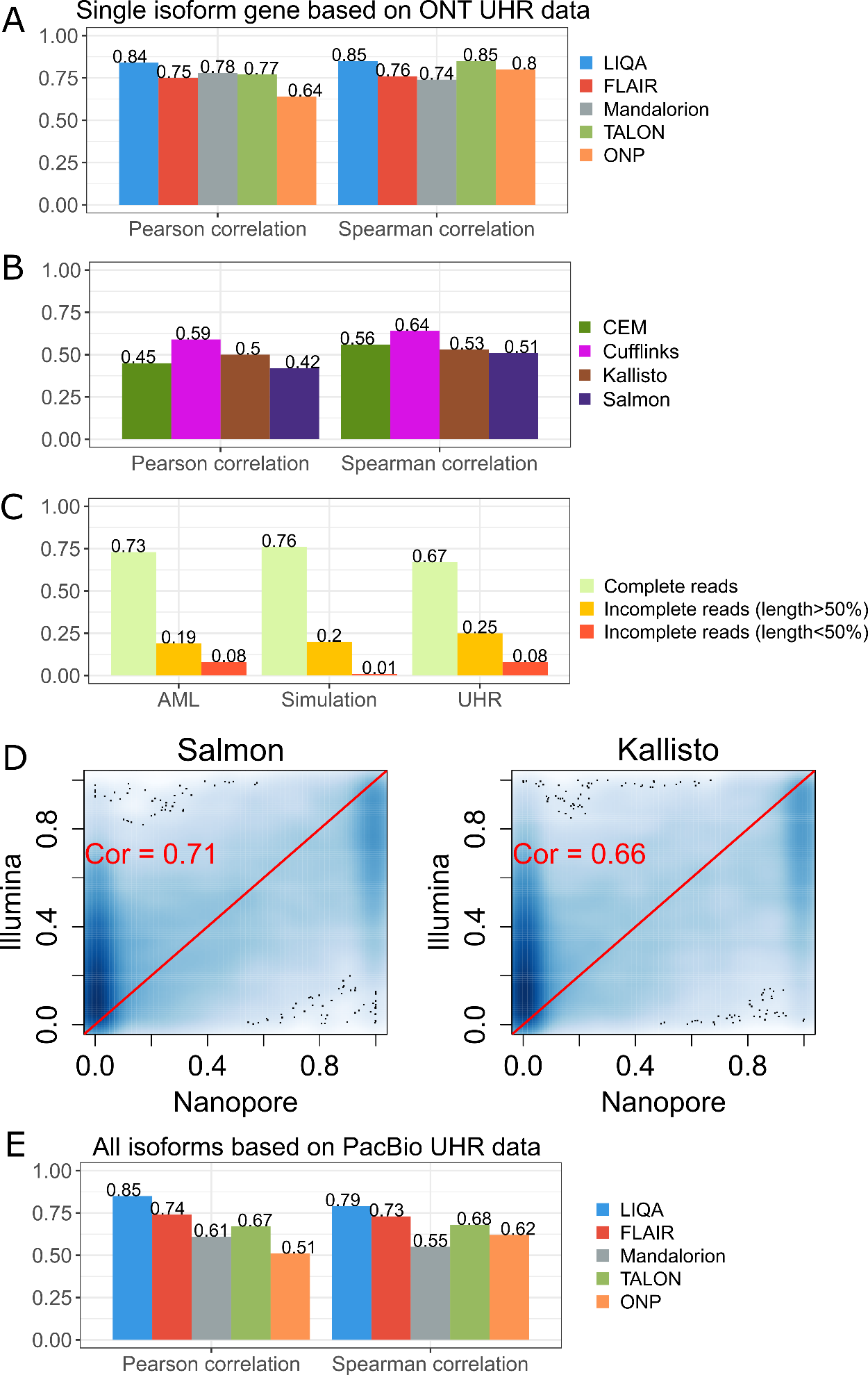

**Supplementary Figure 8. Analysis results of LIQA using real datasets** (A) for the UHR data generated by Nanopore direct mRNA sequencing: Pearson correlation coefficients (left) and spearman correlation coefficient (right) between estimated isoform-specific expression versus qRT-PCR measurements in log scale based on single-isoform genes. (B) for the UHR data generated by Nanopore direct mRNA sequencing: Pearson correlation coefficients (left) and spearman correlation coefficient (right) between estimated isoform-specific expression versus qRT-PCR measurements in log scale using short-read methods (CEM, Cufflinks, Kallisto, Salmon). (C) Distribution of complete and incomplete reads from three datasets (AML, Simulation, UHR by Nanopore). (D) Scatter plots of estimated isoform relative abundances using long-read data (LIQA) versus short-read data (Salmon and Kallisto) from the AML sample. (E) Analysis on another PacBio-based UHR dataset with increasing sequencing depth. Pearson correlation coefficients (left) and spearman correlation coefficient (right) between estimated isoform-specific expression versus qRT-PCR measurements in log scale.

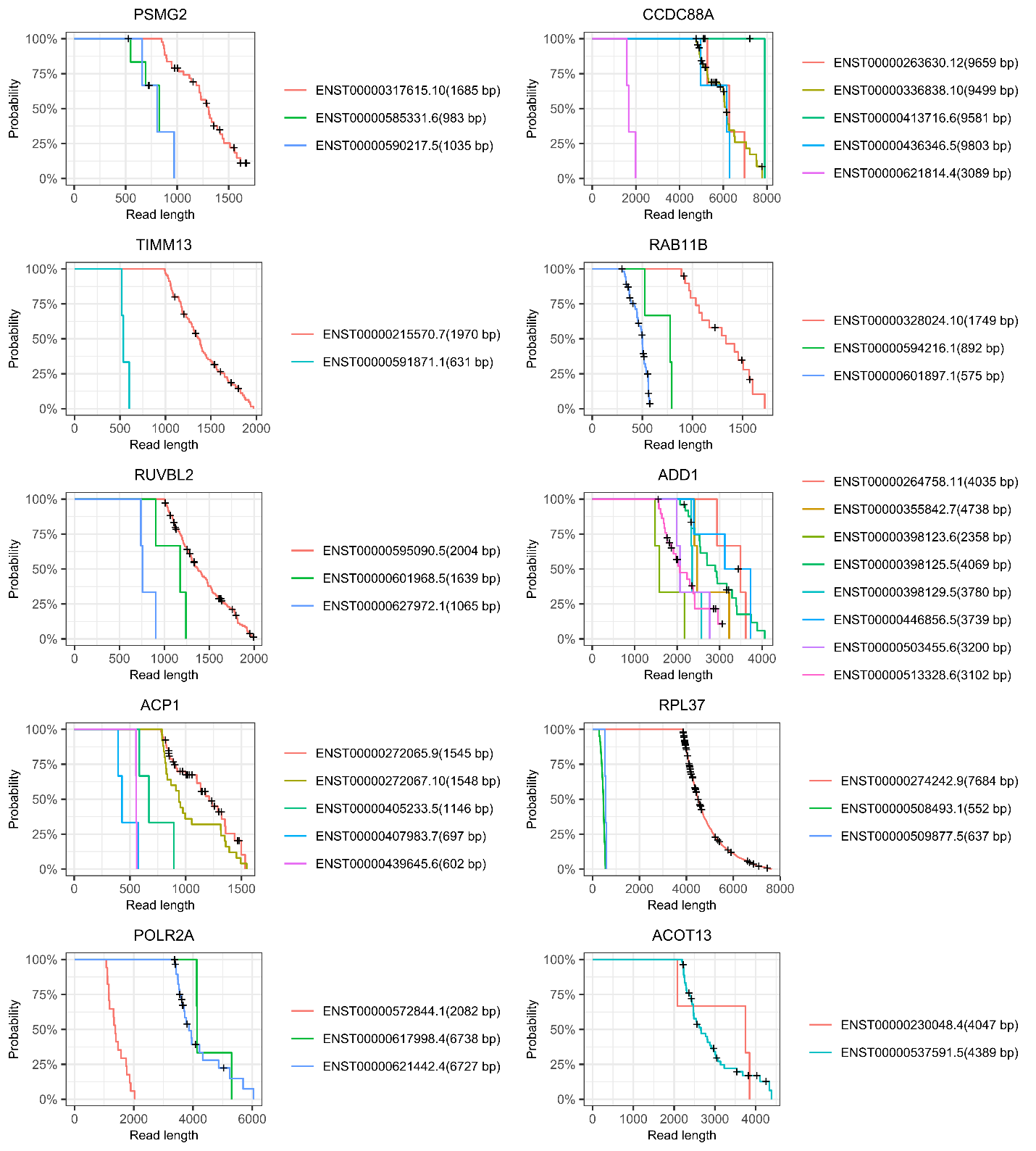

**Supplementary Figure 9.** **Isoform-specific read length probability estimations based on simulated data.** Isoform-specific read length probability ($f(L_{r}>l)$) from 9 randomly selected genes selected were calculated based on simulated data.

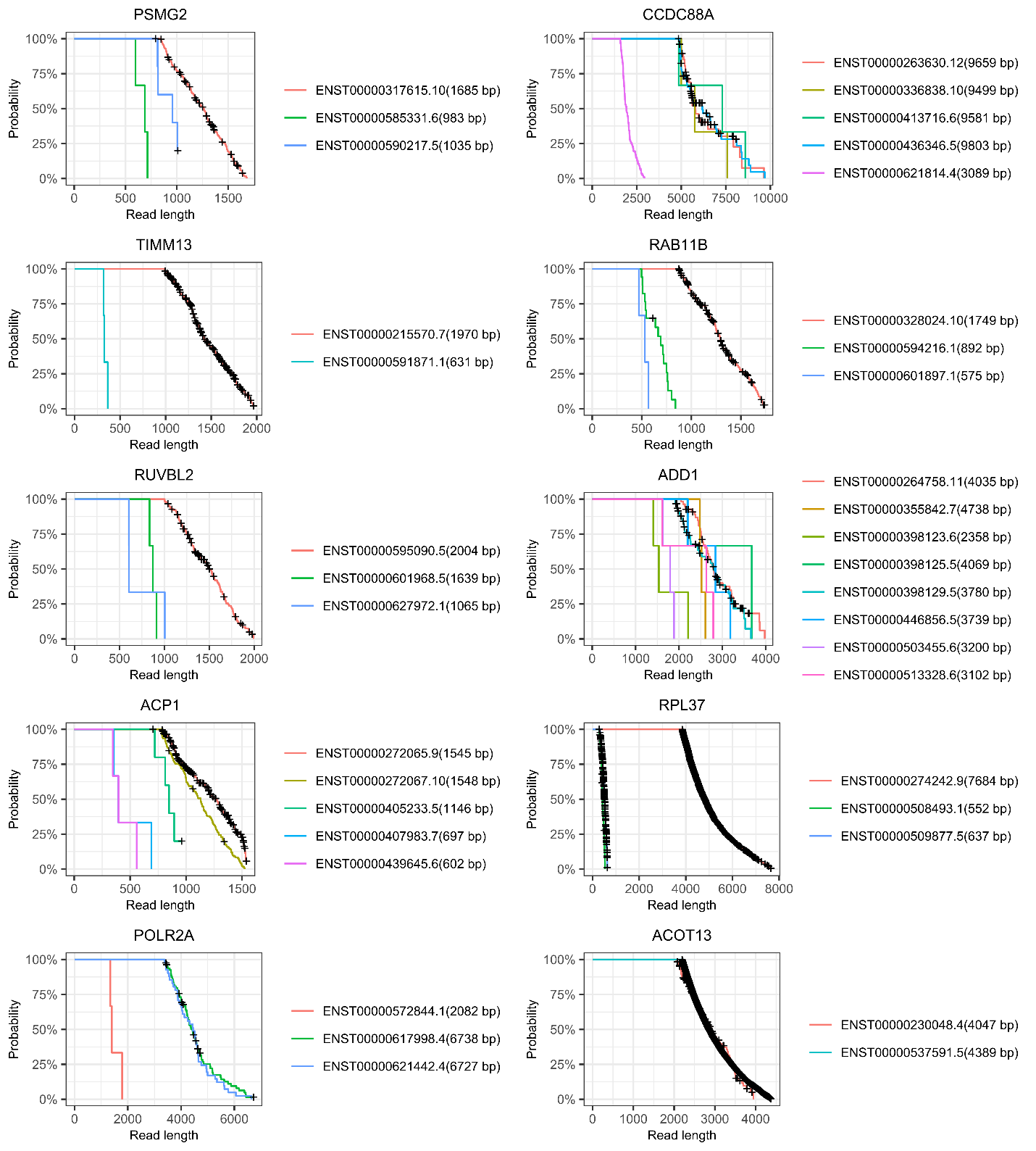

**Supplementary Figure 10.** **Isoform-specific read length probability estimations based on UHR data.** Isoform-specific read length probability ($f(L_{r}>l)$) from 9 randomly selected genes selected were calculated based on UHR data.

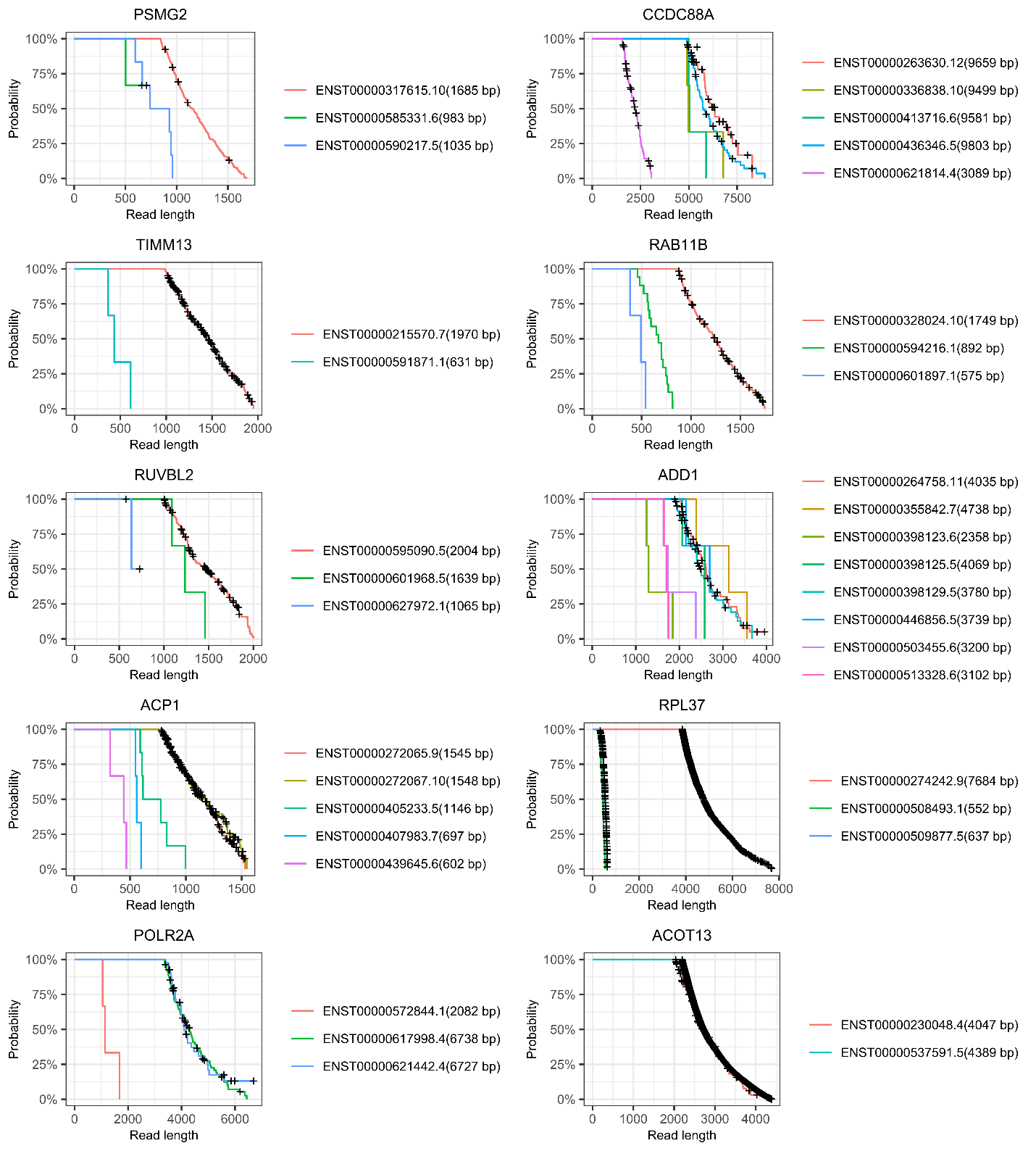

**Supplementary Figure 11.** **Isoform-specific read length probability estimations based on AML data.** Isoform-specific read length probability ($f(L_{r}>l)$) from 9 randomly selected genes selected were calculated based on AML data.

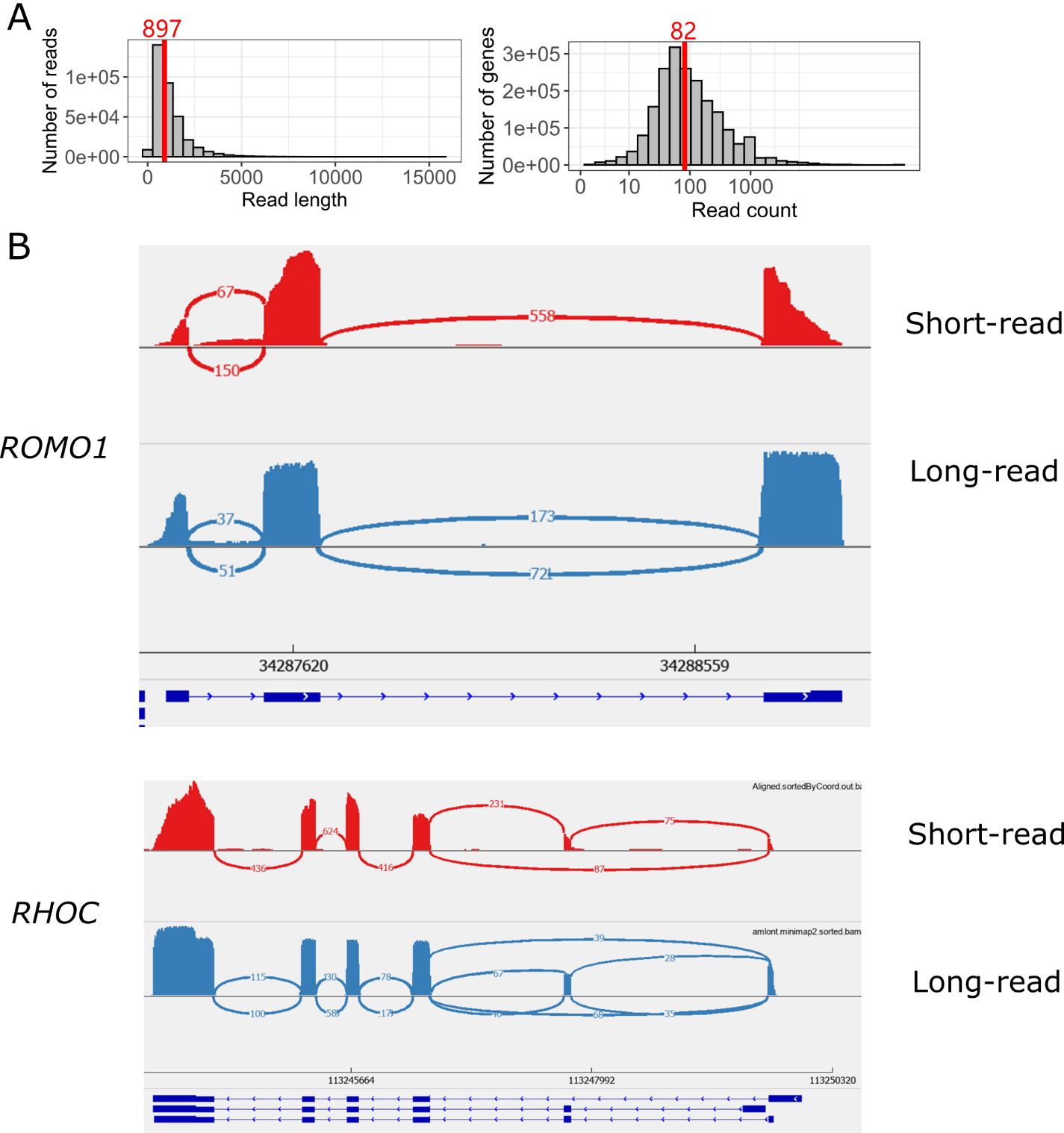

**Supplementary Figure 12. Summary of the Nanopore direct mRNA sequencing data on the UHR sample.** (A) Characteristics of UHR data. Read length distribution (left) and read count distribution by genes in log-scale (right) (B) Examination of read coverage difference between Illumina and Nanopore data on 2 randomly selected genes.

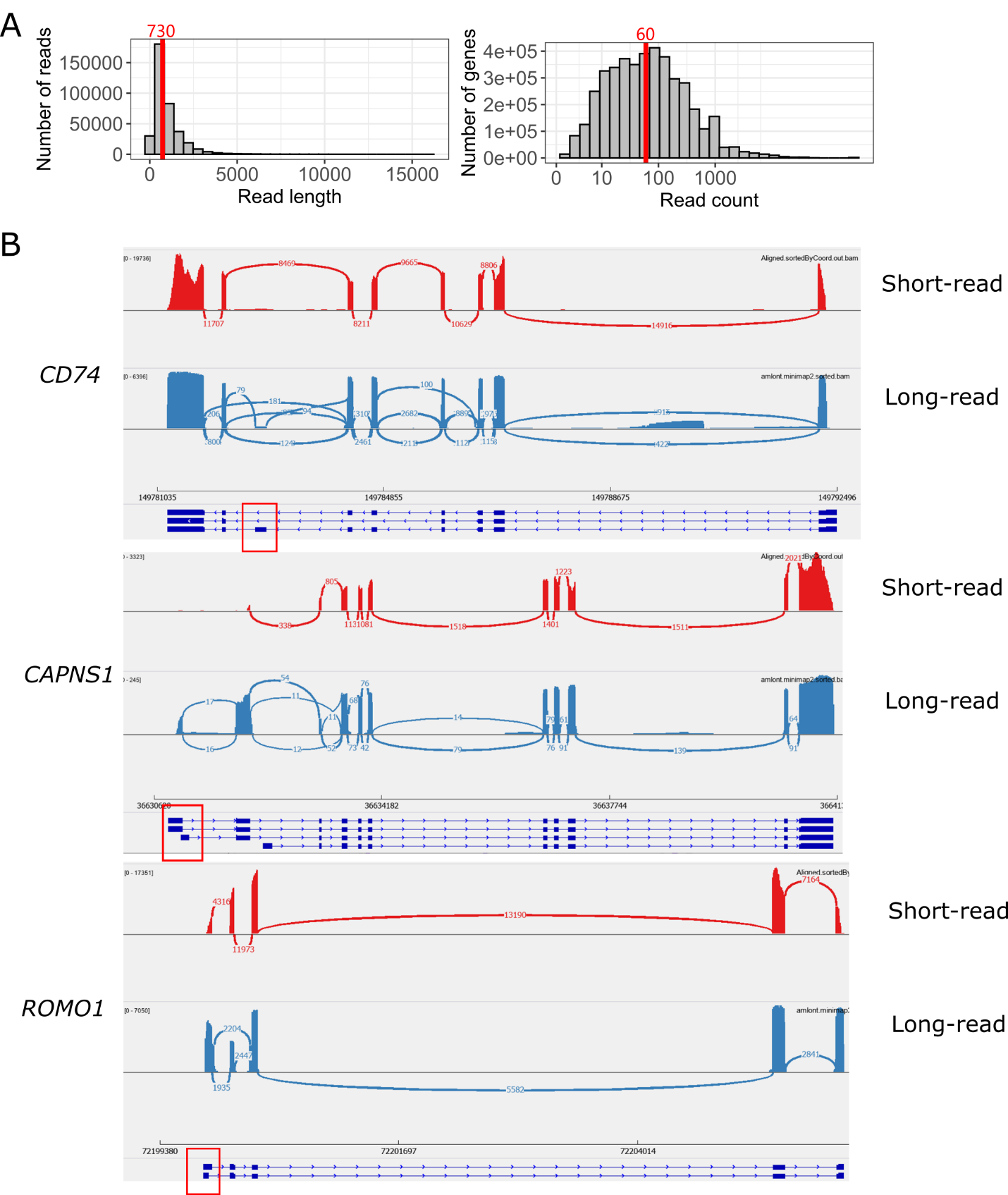

**Supplementary Figure 13. Summary of the Nanopore cDNA sequencing data on the AML sample.** (A) Characteristics of AML data. Read length distribution (left) and read count distribution by genes in log-scale (right) (B) Examination of isoform usage inferred by LIQA. Sashimi plots of gene *CD74, CAPNS1* and *ROMO1*. Informative exonic regions were in red squares.

**Supplementary Tables**

| **Method/Read coverage** | **Runtime (Min)** | | | |
| --- | --- | --- | --- | --- |
|  | **0.5M** | **1M** | **3M** | **5M** |
| LIQA | 43 | 49 | 55 | 63 |
| FLAIR | 21 | 23 | 26 | 35 |
| Mandalorion | 33 | 37 | 41 | 42 |
| TALON | 19 | 25 | 27 | 28 |
| ONP | 8 | 10 | 10 | 11 |

**Supplementary Table 1. Runtime comparison between different methods on simulation data.**

| **Statistical comparison of spearman correlation of TPM** | | | | | |
| --- | --- | --- | --- | --- | --- |
| **Read coverage** | **Method** | LIQA | FLAIR | Mandalorion | TALON |
| 5M | FLAIR | 0.67(0.02,0.05) | NA | NA | NA |
|  | Mandalorion | 0.016(0.18,0.07) | 0.036(0.16,0.08) | NA | NA |
|  | TALON | 0.197(0.05,0.04) | 0.467(0.03,0.04) | 0.069(0.13,0.07) | NA |
|  | ONP | <0.001(0.16,0.05) | 0.004(0.14,0.05) | 0.791(0.02,0.08) | 0.006(0.11,0.04) |
| 3M | FLAIR | 0.118(0.07,0.04) | NA | NA | NA |
|  | Mandalorion | 0.052(0.14,0.07) | 0.341(0.07,0.07) | NA | NA |
|  | TALON | 0.016(0.09,0.04) | 0.617(0.02,0.04) | 0.47(0.05,0.07) | NA |
|  | ONP | 0.004(0.13,0.04) | 0.201(0.06,0.05) | 0.892(0.01,0.07) | 0.317(0.04,0.04) |
| 1M | FLAIR | 0.108(0.07,0.04) | NA | NA | NA |
|  | Mandalorion | <0.001(0.23,0.07) | 0.025(0.16,0.07) | NA | NA |
|  | TALON | 0.005(0.1,0.04) | 0.448(0.03,0.04) | 0.052(0.13,0.07) | NA |
|  | ONP | <0.001(0.15,0.04) | 0.081(0.08,0.05) | 0.258(0.08,0.07) | 0.191(0.05,0.04) |
| 0.5M | FLAIR | 0.033(0.1,0.05) | NA | NA | NA |
|  | Mandalorion | <0.001(0.29,0.07) | 0.013(0.19,0.08) | NA | NA |
|  | TALON | <0.001(0.19,0.04) | 0.029(0.09,0.04) | 0.161(0.1,0.07) | NA |
|  | ONP | <0.001(0.3,0.05) | <0.001(0.2,0.05) | 0.895(0.01,0.08) | 0.006(0.11,0.04) |

**Supplementary Table 2. Pairwise statistical comparison of spearman correlation of TPM.** In each cell, p-value (mean difference, standard deviation of the difference) are calculated. Significant comparisons are highlighted in red at 0.05 significance level while others are grey.

| **Statistical comparison of spearman correlation of relative abundance** | | | | | |
| --- | --- | --- | --- | --- | --- |
| **Read coverage** | **Method** | LIQA | FLAIR | Mandalorion | TALON |
| 5M | FLAIR | 0.653(0.02,0.04) | NA | NA | NA |
|  | Mandalorion | 0.011(0.18,0.07) | 0.027(0.16,0.07) | NA | NA |
|  | TALON | 0.174(0.05,0.04) | 0.443(0.03,0.04) | 0.055(0.13,0.07) | NA |
|  | ONP | <0.001(0.16,0.04) | 0.002(0.14,0.05) | 0.78(0.02,0.07) | 0.004(0.11,0.04) |
| 3M | FLAIR | 0.099(0.07,0.04) | NA | NA | NA |
|  | Mandalorion | 0.041(0.14,0.07) | 0.315(0.07,0.07) | NA | NA |
|  | TALON | 0.011(0.09,0.04) | 0.598(0.02,0.04) | 0.447(0.05,0.07) | NA |
|  | ONP | 0.002(0.13,0.04) | 0.178(0.06,0.04) | 0.886(0.01,0.07) | 0.292(0.04,0.04) |
| 1M | FLAIR | 0.09(0.07,0.04) | NA | NA | NA |
|  | Mandalorion | <0.001(0.23,0.07) | 0.018(0.16,0.07) | NA | NA |
|  | TALON | 0.003(0.1,0.03) | 0.423(0.03,0.04) | 0.04(0.13,0.06) | NA |
|  | ONP | <0.001(0.15,0.04) | 0.066(0.08,0.04) | 0.233(0.08,0.07) | 0.168(0.05,0.04) |
| 0.5M | FLAIR | 0.025(0.1,0.04) | NA | NA | NA |
|  | Mandalorion | 0.007(0.19,0.07) | 0.213(0.09,0.07) | NA | NA |
|  | TALON | 0.014(0.09,0.04) | 0.798(0.01,0.04) | 0.14(0.1,0.07) | NA |
|  | ONP | <0.001(0.2,0.04) | 0.028(0.1,0.05) | 0.889(0.01,0.07) | 0.004(0.11,0.04) |

**Supplementary Table 3. Pairwise statistical comparison of spearman correlation of relative abundance.** In each cell, p-value (mean difference, standard deviation of the difference) are calculated. Significant comparisons are highlighted in red at 0.05 significance level while others are grey.

| **Statistical comparison of spearman correlation of TPM (0.5M dataset)** | | | | | |
| --- | --- | --- | --- | --- | --- |
|  | **Method** | LIQA | FLAIR | Mandalorion | TALON |
| 5' terminal exon | FLAIR | 0.025(0.1,0.04) | NA | NA | NA |
|  | Mandalorion | 0.007(0.19,0.07) | 0.213(0.09,0.07) | NA | NA |
|  | TALON | 0.014(0.09,0.04) | 0.798(0.01,0.04) | 0.14(0.1,0.07) | NA |
|  | ONP | <0.001(0.2,0.04) | 0.028(0.1,0.05) | 0.889(0.01,0.07) | 0.004(0.11,0.04) |
| 3' terminal exon | FLAIR | 0.637(0.02,0.04) | NA | NA | NA |
|  | Mandalorion | 0.019(0.16,0.07) | 0.045(0.14,0.07) | NA | NA |
|  | TALON | 0.091(0.06,0.04) | 0.292(0.04,0.04) | 0.128(0.1,0.07) | NA |
|  | ONP | <0.001(0.15,0.04) | 0.003(0.13,0.04) | 0.886(0.01,0.07) | 0.018(0.09,0.04) |
| chisq>median | FLAIR | 0.167(0.06,0.04) | NA | NA | NA |
|  | Mandalorion | 0.043(0.14,0.07) | 0.256(0.08,0.07) | NA | NA |
|  | TALON | 0.005(0.1,0.04) | 0.294(0.04,0.04) | 0.545(0.04,0.07) | NA |
|  | ONP | <0.001(0.15,0.04) | 0.042(0.09,0.04) | 0.886(0.01,0.07) | 0.176(0.05,0.04) |
| chisq<median | FLAIR | 0.809(0.01,0.04) | NA | NA | NA |
|  | Mandalorion | 0.036(0.14,0.07) | 0.056(0.13,0.07) | NA | NA |
|  | TALON | 0.248(0.04,0.03) | 0.417(0.03,0.04) | 0.119(0.1,0.06) | NA |
|  | ONP | 0.004(0.12,0.04) | 0.011(0.11,0.04) | 0.768(0.02,0.07) | 0.031(0.08,0.04) |

**Supplementary Table 4. Pairwise statistical comparison of spearman correlation of TPM at different exons or isoforms.** In each cell, p-value (mean difference, standard deviation of the difference) are calculated. Significant comparisons are highlighted in red at 0.05 significance level while others are grey.

| **Statistical comparison of spearman correlation of TPM (gene-level)** | | | | | |
| --- | --- | --- | --- | --- | --- |
| **Read coverage** | **Method** | LIQA | FLAIR | Mandalorion | TALON |
| 5M | FLAIR | 0.634(0.03,0.06) | NA | NA | NA |
|  | Mandalorion | 0.426(0.08,0.1) | 0.282(0.11,0.1) | NA | NA |
|  | TALON | 0.7(0.02,0.05) | 0.366(0.05,0.06) | 0.531(0.06,0.1) | NA |
|  | ONP | 0.329(0.06,0.06) | 0.162(0.09,0.06) | 0.843(0.02,0.1) | 0.456(0.04,0.05) |
| 3M | FLAIR | 0.067(0.11,0.06) | NA | NA | NA |
|  | Mandalorion | 0.408(0.08,0.1) | 0.761(0.03,0.1) | NA | NA |
|  | TALON | 0.046(0.1,0.05) | 0.852(0.01,0.05) | 0.83(0.02,0.09) | NA |
|  | ONP | 0.067(0.11,0.06) | 1(0,0.06) | 0.761(0.03,0.1) | 0.852(0.01,0.05) |
| 1M | FLAIR | 0.254(0.07,0.06) | NA | NA | NA |
|  | Mandalorion | 0.474(0.07,0.1) | 1(0,0.1) | NA | NA |
|  | TALON | 0.167(0.07,0.05) | 1(0,0.05) | 1(0,0.09) | NA |
|  | ONP | 0.066(0.11,0.06) | 0.524(0.04,0.06) | 0.685(0.04,0.1) | 0.444(0.04,0.05) |
| 0.5M | FLAIR | 0.305(0.06,0.06) | NA | NA | NA |
|  | Mandalorion | 0.671(0.04,0.09) | 0.835(0.02,0.1) | NA | NA |
|  | TALON | 0.22(0.06,0.05) | 0.022(0.12,0.05) | 0.27(0.1,0.09) | NA |
|  | ONP | 0.732(0.02,0.06) | 0.514(0.04,0.06) | 0.835(0.02,0.1) | 0.126(0.08,0.05) |

**Supplementary Table 5. Pairwise statistical comparison of spearman correlation of TPM at gene level.** In each cell, p-value (mean difference, standard deviation of the difference) are calculated. Significant comparisons are highlighted in red at 0.05 significance level while others are grey.

| **Statistical comparison of spearman correlation of TPM** | | | | |
| --- | --- | --- | --- | --- |
|  | **Model** | Full model | Read length model | Read quality model |
| Chisq > median | Read length model | 0.451(0.03,0.04) | NA | NA |
|  | Read quality model | 0.156(0.09,0.06) | 0.353(0.06,0.06) | NA |
|  | Baseline model | <0.001(0.13,0.03) | 0.004(0.1,0.03) | 0.509(0.04,0.06) |
| Read quality < median | Read length model | 0.151(0.06,0.04) | NA | NA |
|  | Read quality model | 0.133(0.1,0.07) | 0.555(0.04,0.07) | NA |
|  | Baseline model | <0.001(0.16,0.03) | 0.006(0.1,0.04) | 0.345(0.06,0.06) |

**Supplementary Table 6. Pairwise statistical comparison of spearman correlation of TPM across different models.** In each cell, p-value (mean difference, standard deviation of the difference) are calculated. Significant comparisons are highlighted in red at 0.05 significance level while others are grey.

| **Model** | **Model comparison** | | |
| --- | --- | --- | --- |
|  | **Degree of freedom** | **Δ(-2logLikelihood)** | **P-value** |
| Read quality model | 156 | 215.13 | 0.012 |
| Read length model | 33 | 90.56 | <0.001 |
| Full model | 189 | 307.97 | <0.001 |

**Supplementary Table 7. Model comparison.** Statistical comparison between different models (full model, read length model, read quality model). The reference control model is baseline model with no bias correction.
